# Dietary zinc limitation dictates lifespan and reproduction trade-offs of *Drosophila* mothers

**DOI:** 10.1101/2024.08.28.610171

**Authors:** Sweta Sarmah, Hannah Thi-Hong Hanh Truong, Gawain McColl, Richard Burke, Christen K Mirth, Matthew DW Piper

## Abstract

Dietary metal ions significantly influence the lifespan and reproduction of *Drosophila* females. In this study, we show that while the depletion of all metal ions from the diet adversely affects reproduction and lifespan, the absence of Zn alone negatively impacts reproduction without adversely altering maternal lifespan, indicating it can dictate resource reallocation between key fitness traits. Although our data show that flies sense varying dietary Zn levels, they apparently do not alter their feeding preferences to optimise egg production when faced with a choice between Zn- depleted or Zn- sufficient food, but they can choose to preferentially oviposit on Zn containing food, perhaps indicating a strategy to assure offspring survival. We also uncovered a role for the *white* gene in sustaining high levels of egg viability when Zn is diluted in the diet. These insights into the role of dietary metal ions, particularly Zn, point to a central role for these dietary micronutrients to indicate environmental quality and so govern trade-offs between lifespan and reproduction in flies.

## Introduction

A balanced diet is a central determinant of health, driving changes in growth, reproduction, stress responses, and ageing (Raubenheimer 2016) (Piper 2008). Exactly how dietary nutrient balance is connected to these traits is not fully known, but there is ample evidence to show that excesses or insufficiencies of specific nutrients can be detrimental (e.g. (Raubenheimer 2005) (Anderson, Raubenheimer et al. 2020) (Simpson 2015) (Simpson 2012)). Because nutrient balancing is so important for maximising evolutionary fitness, animals have evolved a wide range of behavioural and physiological adaptations to match nutrient acquisition and metabolism to the available supply (Simpson 2012).

Heterotrophic organisms require a mix of both macronutrients (proteins and carbohydrates) and micronutrients (metal ions, vitamins, and sterols) in their diets (Sang 1961) (Piper 2014) (Mirth 2019). Unlike macronutrients, which are required in relatively large amounts to build biological structures and for use as a source of energy, micronutrients are required in smaller amounts, and are mostly required to facilitate metabolic processes (e.g. as enzyme co-factors) and to maintain the physical environment (Bhattacharya 2016). Because of their relative abundance and their role in energy generation, most research on the eco-evolutionary significance of dietary nutrition has focused on macronutrients, with relatively less emphasis on micronutrients.

In *Drosophila melanogaster*, and other insects, several studies have shown that increasing the ratio of dietary protein relative to carbohydrate can promote adult reproduction, but at the cost of reduced lifespan, while diets with lower protein to carbohydrate proportions reduce reproduction and increase lifespan (Mair 2005) (Piper 2011) (Simpson 2017) (Lee 2008) (Skorupa 2008). More recent data has shown that the reciprocal relationship between these traits is conditional on the quality of protein provided and the levels of sterol in the food: for female *D. melanogaster*, a combination of small amounts of high quality protein with ample amounts of the micronutrient cholesterol provides sustenance for both long lifespan and high reproduction simultaneously (Piper 2017) (Zanco 2021). These data show that micronutrients can mediate the way that macronutrients affect fitness.

We have also found that manipulating the levels of some metal ions, which are also essential micronutrients for development, plays a central role in determining fitness of female flies by controlling rates of reproduction and their relationship to lifespan (Piper 2014). These data indicate that flies sense the levels of one or more metal ions in the process of determining the degree to which they commit resources to reproduction versus adult survival. Despite this apparent central role in life history trait trade-offs, the scope of fitness responses to dietary metal ions remains under-studied.

Metal ions are important structural and catalytic components that support a wide range of processes that collectively contribute to an organism’s overall well-being. For example, iron (Fe), copper (Cu), magnesium (Mg), manganese (Mn) and Zinc (Zn) all function as catalysts in various enzymes involved in energy generation, amino acid metabolism, and cellular redox metabolism (Jomova, 2022) (Dow, 2017). Additionally, Zn is integral to the structural integrity and function of proteins engaged in DNA repair, transcriptional regulation, and translational processes, while iron is involved in oxidative stress mitigation, and copper is a vital cofactor for enzymes participating in respiration and pigmentation (Vilas 2018) (Galaris 2019) (Tsang 2021). These molecular functions underpin important physiological processes and depletion of these metal ions has been linked to growth inhibition, sterility, loss of circadian rhythms and immunodeficiency (Hu 2020) (Sang 1961) (Tian 2014) (Iatsenko 2020) (Rudisill 2019) (Kim 2010) (Tian 2012) (Que 2015) (Tian 2014) (Horner 2008) (Pankau 2022).

Here, we explore the effects of dietary metals on adult female egg production, behaviour, and lifespan in *D. melanogaster*. Interestingly, while depriving flies of all of the metal ions generally reduces reproduction and shortens lifespan, manipulating only dietary Zn results in a lifespan / reproduction trade-off similar to that observed when dietary protein: carbohydrate ratios are altered. These data indicate that flies have physiological sensing mechanisms that enable them to measure dietary Zn supply and adjust their life history strategies accordingly. These observations offer important insights into how dietary components, in particular the micronutrients, shape evolutionary fitness.

## Results

### Dietary metal ions are required for lifespan and reproduction in *Drosophila* females

To examine the effects of dietary metal ions on *Drosophila* lifespan and fecundity, we used completely defined, synthetic (holidic) diets (Piper 2014) (Piper 2017). Four different experimental diets were created by diluting the mixture of metal ions (Ca, Cu, Fe, Mg, Mn, and Zn) to 0%, 10%, 50% and 100% of the level in the complete (control) diet, while all other components were identical between foods (Figure S1A). We tested fecundity and lifespan responses to these treatments using two outbred strains of *Drosophila* (Dahomey): one wild-type (red-eyed), and the other carrying a mutation at the *white* locus (white-eyed). The *white* gene is known to encode a pigment transporter, which is important for eye colour, and has also been shown to play a role in whole body metal ion homeostasis (Tejeda-Guzmán 2018).

Both red-eyed Dahomey (*rDah*) and white-eyed Dahomey (*wDah*) suffered progressively shortened lifespan as metal ions were diluted below 50% of the level on control food to 10% and 0%, but they did so to significantly different extents (Figure 1A-B, Table S1). In particular, *rDah* flies showed a greater lifespan shortening at 10% and 0% metal ions (medians: ∼14 days and ∼9 days respectively) than *wDah* (medians: ∼23 days and ∼13 days respectively; Figure 1C-D, Table S2).

**Figure 1.**
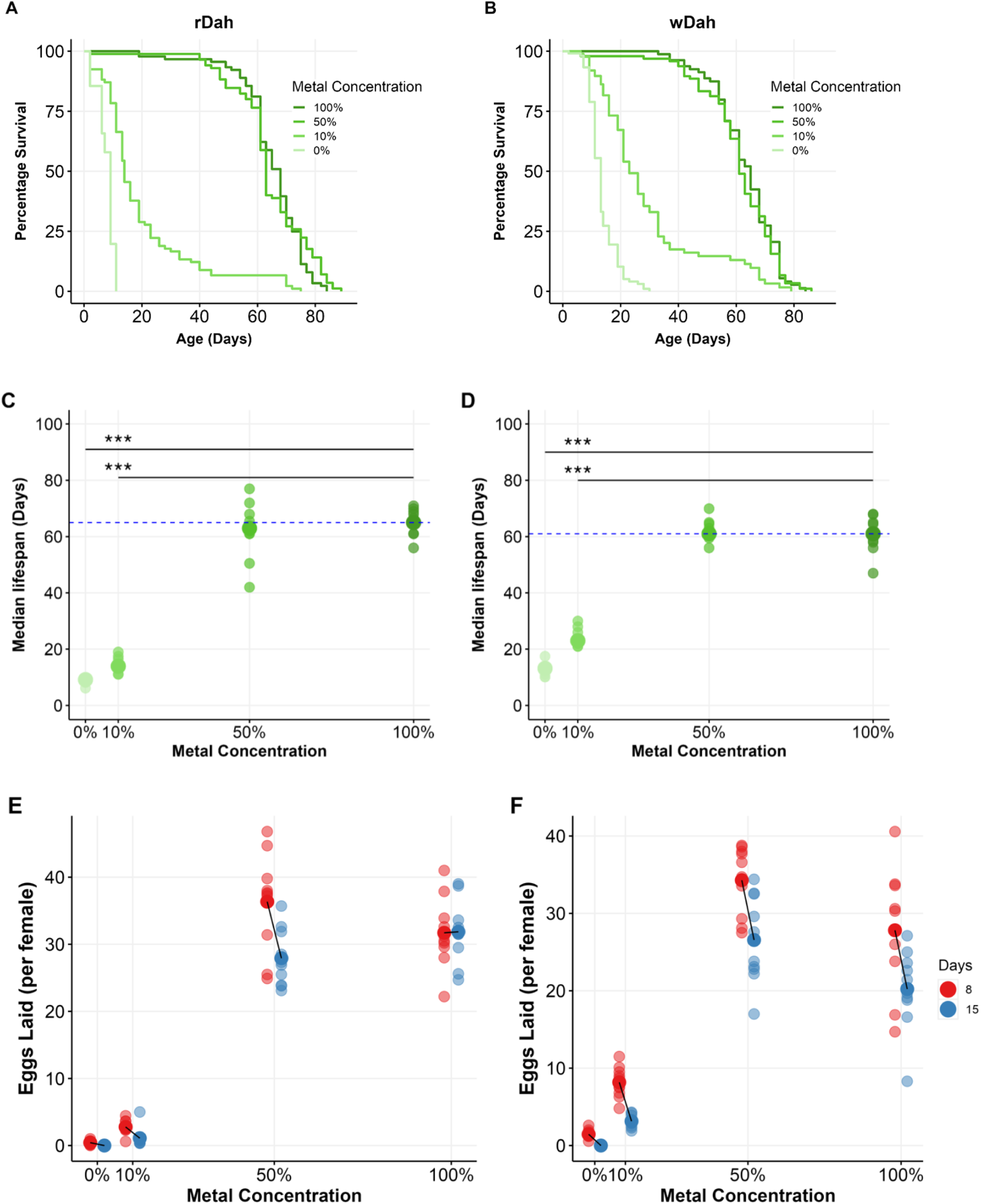
Diluting dietary metal ions reduces the lifespan and fecundity of Drosophila. (A) rDah and (B) wDah females demonstrate a significantly reduced lifespan with metal ion dilution below 50% of the level in control food. (C) and (D) Replicate (small circle) and median (large circle) lifespans for data illustrated in panels (A) and (B), respectively. The horizontal dashed blue line denotes the median lifespan of the ‘100%’ metal ion (fully fed control) group. The asterisks indicate significant difference from 100% metal ions (Table S2). (E) rDah females and (F) wDah females display a similar, but statistically different (Table S3), decline in egg laying over time and in response to metal ion dilution. Asterisks indicate significant lifespan differences from the 100% metal ion control group (***, p < 0.001; see Table S2 for details).

Fecundity, and its decline with fly age, were also modified by metal ion dilution, and these effects were also altered by the flies’ genotype (Figure 1E-F; Table S3). Specifically, the egg production of flies fed 0% and 10% metal ions was significantly reduced from that of flies on 100% control food, and this reduction was more pronounced in *rDah* than *wDah* flies. These findings demonstrate the essential nature of dietary metal ions to *Drosophila* adult females for egg laying and lifespan. They also show that the *white* transporter has a role to play in egg production and lifespan during metal ion limitation.

### Genotype modifies the lifespan and fecundity responses to individual metal ion restrictions

Since metal ions are essential for egg production and adult lifespan, we wondered if the flies suffered equally detrimental outcomes for deprivation of any one metal ion, or if these responses would be distinct, which could indicate differences in their biological and ecological significance. To answer this question, we individually omitted each of the metal ions from the diet and measured fly lifespan and fecundity (Figure 2, Figure S2).

**Figure 2.**
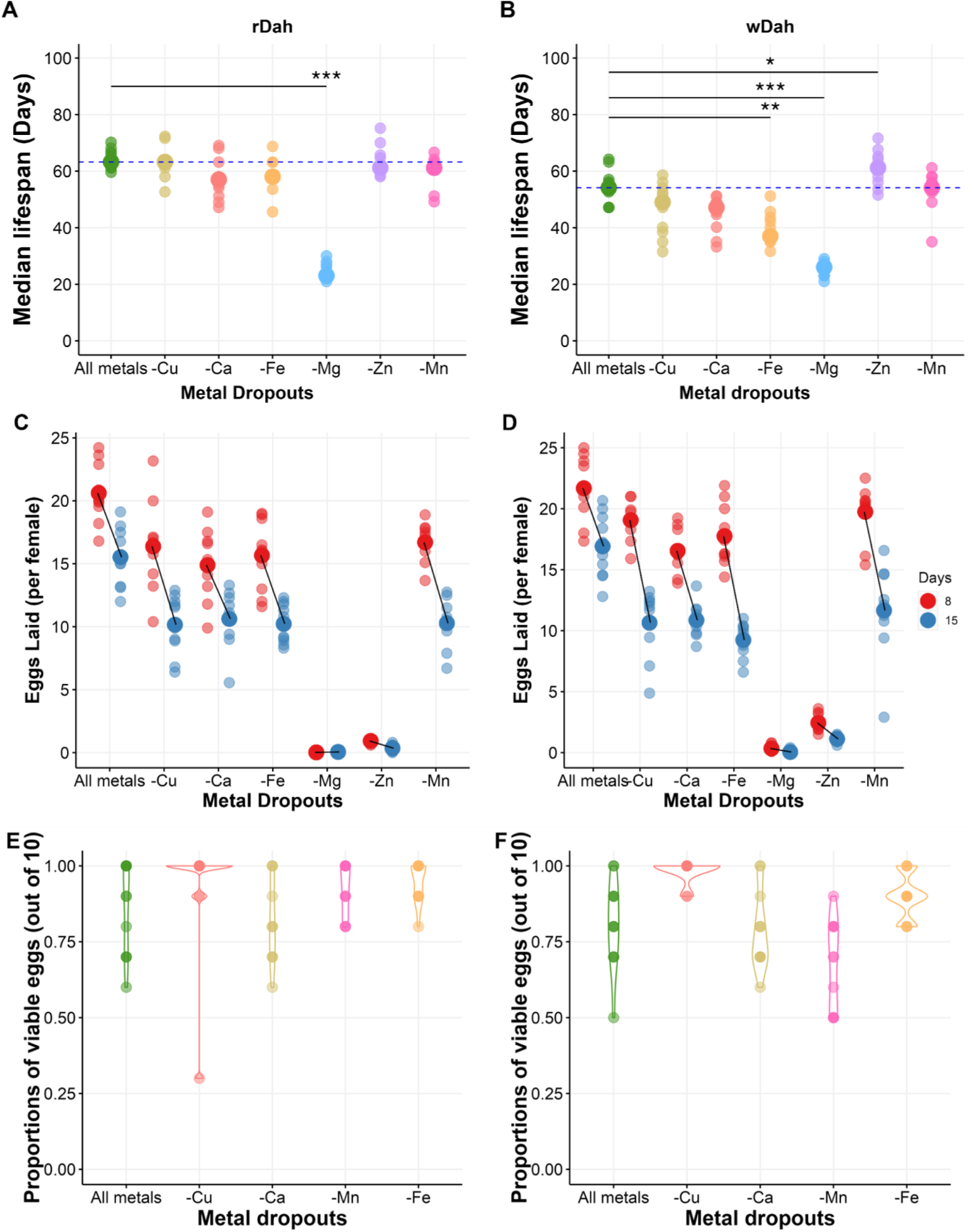
Individual metal ion depletions elicit distinct outcomes for both lifespan and reproduction. For rDah flies (A), deprivation for magnesium resulted in significant lifespan shortening. In contrast, wDah flies (B) exhibited a significant lifespan cost when deprived of Fe or Mg, and a small but significant lifespan benefit when deprived of Zn. Replicate means are represented by small circles, while larger circles represent population median. To facilitate comparisons between treatment groups, a dashed horizontal line marks the mean lifespan of the ‘All metal’ treatment group. The asterisks indicate a significant difference from “All metals”. Full lifespan curves are shown in supplementary Figure S2. After exposure to each diet for 8 or 15 days of adulthood, egg laying was relatively unaffected, except for when flies were exposed to Zn or Mg depleted foods, on which both rDah (C) and wDah (D) showed strongly reduced egg production. For both rDah (E) and wDah (F) egg-to-adult viability was not compromised across different individual metal dropouts (Ca, Cu, Fe and Mn). Asterisks indicate significant lifespan differences from “All metals” control group (***, p < 0.001; **, p < 0.01, *, p < 0.05, see Table S5 for details).

The effects of individual metal ion omissions on lifespan differed according to the identity of the metal ion dropped out, and this effect was modified by genotype (Figure 2A-B, Figure S2 B-C, Table S4). Specifically, there was very little effect of individual metal ion depletion on *rDah* lifespan, except for Mg removal, which shortened median lifespan from ∼62 days to ∼23 days. By contrast, the responses of *wDah* varied more, as Fe, Mg and Zn all significantly modified lifespan to some extent (Figure 2B, Figure S2C). Specifically, *wDah* median lifespan was shortened by Fe and Mg omission, while surprisingly, Zn omssion, caused a mild, but significant, extension to median lifespan (Figure 2B, Figure S2C, Table S5).

Individual metal ion deprivations also elicited distinct outcomes for fecundity, but these effects were not modified by genotype (Figure 2C-D, Table S6). The most striking dietary effects were caused by deprivation of either Mg or Zn, which resulted in an arrest of egg production by day 8 for both genotypes; none of the other metal ion dropouts differed substantially from the pattern of egg laying of fully-fed controls (Figure 2C-D).

When adult female flies encounter food with inadequate levels of Ca, Cu, Fe, or Mn, continuing to lay eggs at a high rate early in life may be an effective strategy to maximise overall fitness as long as the eggs laid are still adequately provisioned with nutrients to be viable. To assess this, we collected eggs from females maintained on each of these single ion deprivation diets for 8 days and transferred the eggs to a yeast-based diet (Bass 2007), which is nutritionally complete, to support larval development. For both *rDah* and *wDah*, deprivation for any one of the metal ion dropouts did not modify the proportion of eggs that produced viable adults (Figure 2E-F, Table S7). Thus, although lacking any one of these metal ions from the adult diet can be costly for maternal lifespan, early fecundity and offspring production remain unaffected by limiting amounts of any one of these nutrients.

Taken together, these data demonstrate that female flies implement at least three different strategies when encountering deprivation for an essential metal ion: (1) they continue to produce and lay viable eggs with either no, or only a relatively small, detrimental effect on maternal lifespan (observed for Cu, Ca, Fe or Mn deprivation); (2) they cease egg production and preserve maternal lifespan (observed for Zn deprivation), or; (3) they cease producing eggs and die young (Mg deprivation).

### Zn dilution reduces reproduction, but maternal lifespan is not compromised

Zn deprivation produced a different outcome than deprivation for any of the other metal ions: the flies rapidly ceased egg production while preserving maternal lifespan. To explore this further, we created four different experimental diets containing Zn at 0%, 10%, 50% and 100% of the level in the complete (control) diet (Figure 3, Figure S3).

**Figure 3.**
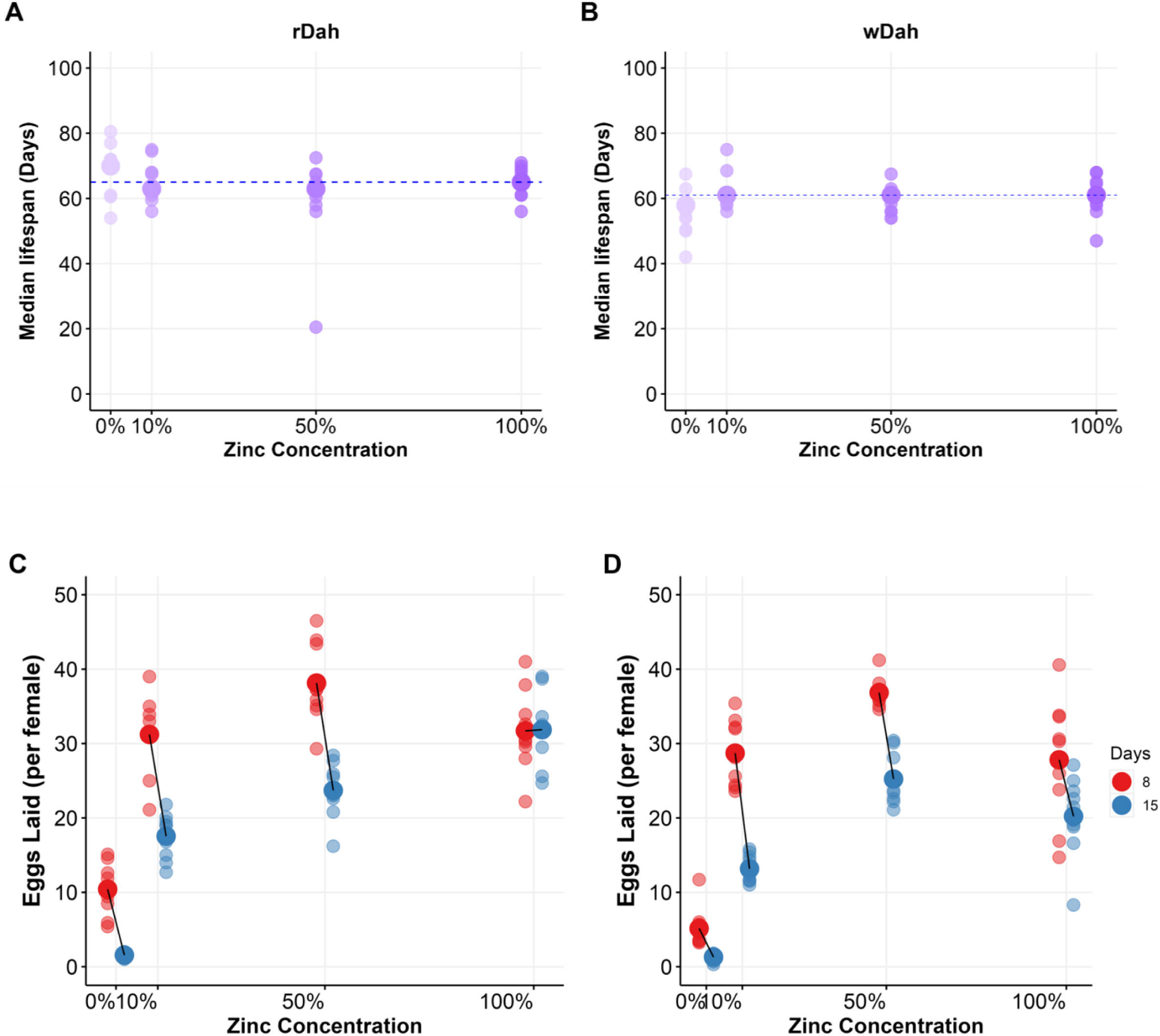
Dietary Zn dilution reduces reproduction with a small increase in lifespan. Dietary Zn dilution had little effect on rDah (A) and wDah (B) lifespan. Population medians are shown as larger circles and smaller circles represent replicates. The dashed horizontal line at the median lifespan value represents the ‘100%’ Zn control group. Full lifespan curves are shown in supplementary Figure S3. Both rDah (C) and wDah (D) females display a significant decline in egg laying over time and in response to Zn dilution. The two genotypes differed significantly in the dynamics of their responses to Zn dilution (Table S10).

There was no reduction in lifespan for either *rDah* or *wDah* flies at any level of Zn dilution (Figure 3A-B, Figure S3B-C, Table S8-9). Furthermore, we did not observe an extension of median lifespan in this experiment, indicating any benefits of Zn restriction to lifespan are dietary context dependent.

Reducing Zn in the diet caused a graded reduction in egg laying that was similar for both genotypes, although the exact pattern of change was significantly modified by genotype (Figure 3C-D, Table S10). Notably, both *wDah* and *rDah* flies exhibited only a small reduction in egg laying with 10% Zn, but a strong reduction with 0% Zn (Figure 3C-D).

One explanation for why flies on Zn depleted diets shut down egg production and maintain adult lifespan is that they simply eat less food. If true, the flies would be eating less protein, meaning that macronoutrient restriction, rather than Zn restriction, might account for the effects we have observed (Mair, 2005) (Lee, 2008) (Skorupa, 2008). To assess this, we measured the change in metal ion levels in Zn depleted flies.

We separated ovaries from the remaining body tissue of *rDah* and *wDah* flies and measured the levels of Zn, Cu, Fe, Mg and Mn using Inductively Coupled Plasma-Mass Spectrometry, comparing the proportions of each metal in flies on a 0 Zn diet with those on a complete diet (100N) (Figure 4). Metal ion depletion was evident in both tissue types in both genotypes, but occurred to different extents for each metal (Table S11). After 8 days of feeding on Zn deficient food, we observed a specific reduction in Zn in the body samples when compared with fully fed flies (Figure 4A-B). This effect was more pronounced in *rDah* (∼5-fold lower) than *wDah* (∼2.5-fold lower) (Table S12). All other metal ions in the bodies of flies feeding on Zn depleted food were either unchanged, or in the case of Cu, exhibited a small but significant increase in abundance (Figure 4A-B, Table S12). This pattern of change indicates that the flies are maintaining the intake of all nutrients, other than Zn for which they are being specifically depleted, which is consistent with the flies not reducing overall food intake on Zn depleted diets.

**Figure 4.**
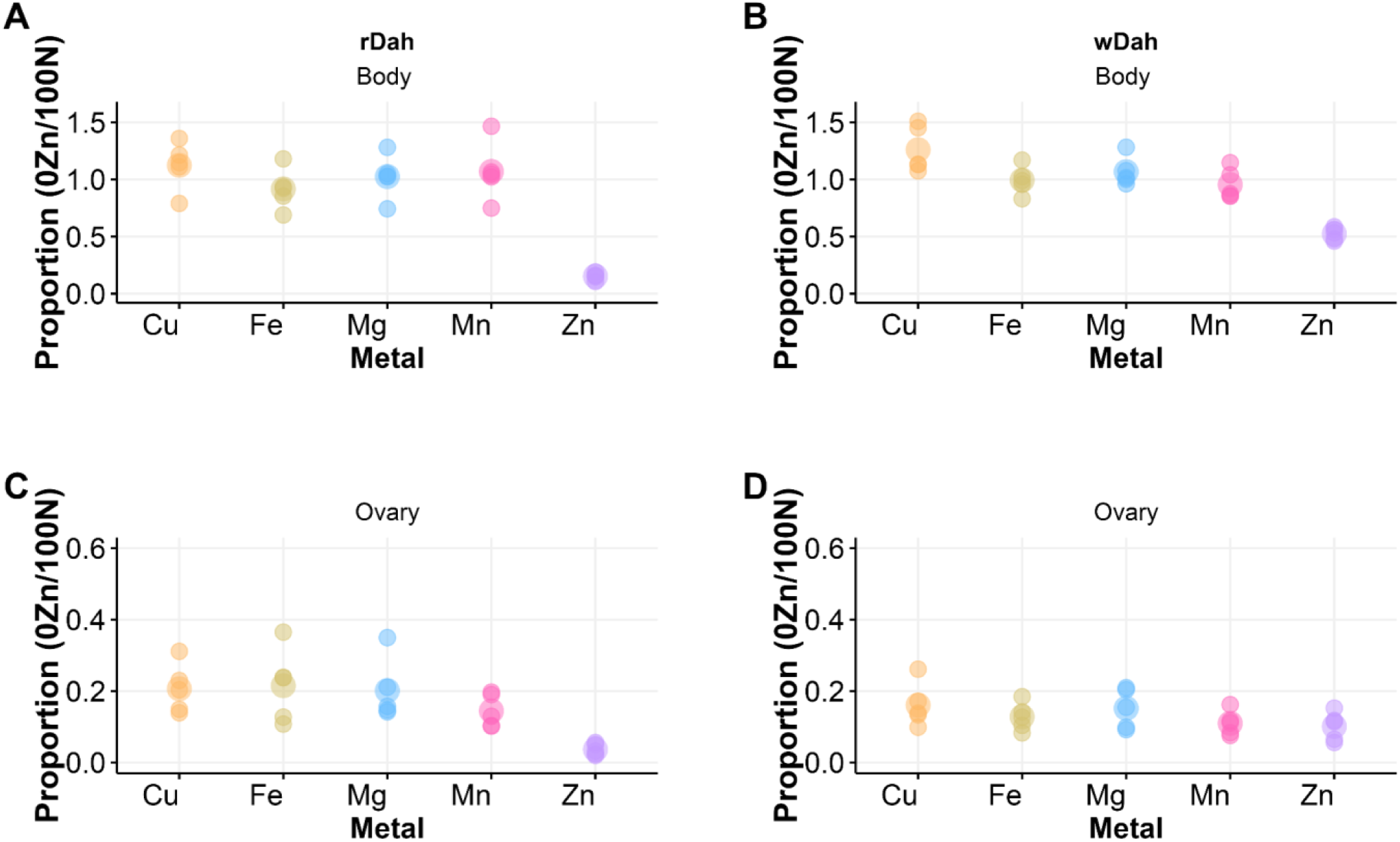
Genotype and diet influence metal ion levels in the ovary and body of flies. Proportions of metal ions in the bodies of (A) rDah and B) wDah flies were significantly modified by dietary Zn levels. Specifically, Zn levels were strongly depleted in the bodies of Zn depleted flies. In the ovary, metal proportions (Cu, Fe, Mg, Mn, Zn) were significantly reduced in flies on Zn depleted diets in both (C) rDah and (D) wDah, perhaps reflecting a reduction in ovary size as Zn depletion suppresses egg production.

In the ovary samples, Zn was also drastically reduced by at least 10-fold when compared to fully fed flies (Figure 4C-D, Table S12). In this case, however, all other metal ions measured also showed ∼5-fold lower levels than what was found in fully fed animals. Given that these flies still retained normal levels of the metal ions in their body tissue (Figure 4A-B), we assume that this is not specific depletion of all metals from the ovaries, but instead reflects a generalised reduction in ovary size that accompanied cessation of egg production (Kosakamoto 2024). Interestingly, although both sets of fly tissues show substantial Zn loss when feeding on Zn depleted diets, the flies still retain enough Zn to sustain vital functions that support full lifespan.

### Although females do not selectively feed on Zn-containing food, they exhibit a preference for oviposition site

Our data show that flies on Zn depleted diets cease egg production and lose a substantial proportion of their body Zn levels. We therefore decided to test if the flies could modify their behaviour to restore fitness when a food choice is available that would allow them to avoid Zn depletion. To do this, we gave groups of females one of several pairwise food choices, in which only the Zn concentration varied, and counted the total number of eggs laid. If dietary Zn levels limit egg production, counting eggs can be used as a proxy for which food the flies choose to eat.

The diet pairs we used were a positive control in which both food options contained 100% Zn (100_100); a negative control in which both foods contained 0% Zn (0_0); and an experimental condition in which the flies could choose between one food containing 0% Zn and another containing 100% Zn (0 _100). To control for the situation where flies randomly sampled the two food options in the experimental condition and so effectively consumed 50% Zn, we added a third control, in which both food options contained 50% Zn (50_50) (Figure S4). For all conditions, we counted eggs laid in a 24-hour window after continuous exposure to the diet pairs for 2 days (Figure S5A-B), 8 days (Figure 5A-B) and 15 days (Figure S5C-D).

**Figure 5.**
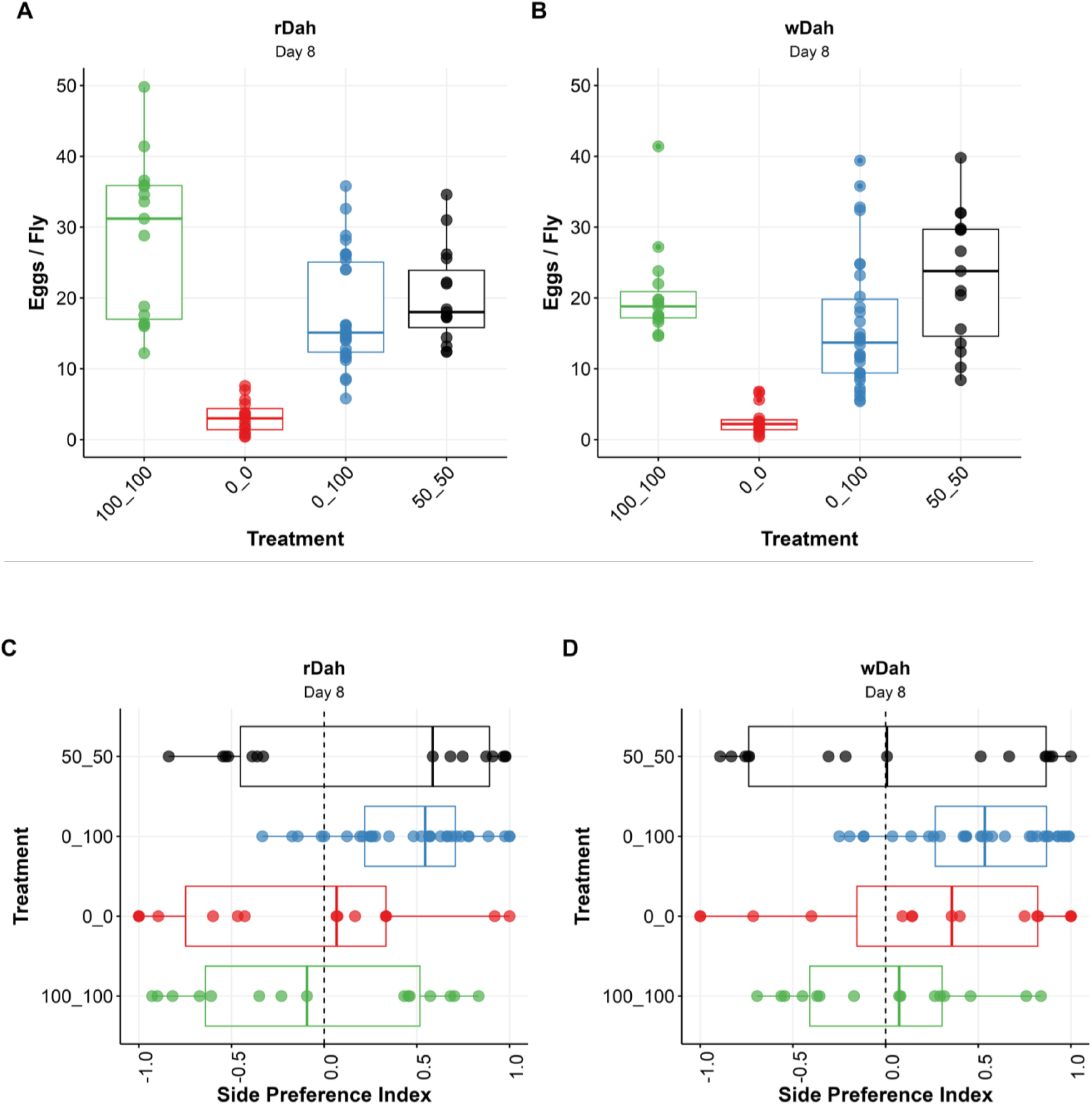
Given pairwise choices of food varying only in Zn content, flies do not optimise egg production, but do select Zn-containing food for oviposition. After 8 days of exposure to a choice of two diets that differed only in Zn concentration, rDah flies (A) and wDah flies (B) laid eggs at a rate that reflected random feeding between the two foods. (C) rDah and (D) wDah flies on day 8 showed a preference to oviposit on food containing Zn over food lacking Zn. This phenotype appeared to strengthen as the assay duration increased (see Figure S5 for day 2 and day 15 data for both assays)(Table S15-16).

For both genotypes, egg production changed in response to the diet choice conditions and the age at which egg laying was assessed (Figure 5A-B, Figure S5A-D, Table S13). At the earliest time point (day 2), egg-laying was mostly unchanged across all dietary pairs indicating that Zn-mediated changes occur more than 2 days after being introduced to the new foods (Figure S5A-B; Table S14).

By day 8 of adulthood, egg production had diverged between diet choice groups (Figure 5A-B, Table S13-14). For both genotypes, egg production was almost completely arrested for flies without Zn (0_0). For *rDah* flies, egg laying significantly increased when flies were maintained in the 50_50 choice, and increased further again for flies confined to the 100_100 choice. These data indicate that egg laying was Zn limited and thus egg production reflected diet choice. When given the experimental diet pair (100_0 choice), egg numbers were higher than for flies in the 0_0 condition, lower than for flies on the 100_100 condition, and no different from the flies that were maintained on the 50_ 50 diet pair (Figure 5A, Table S14). These data are consistent with a situation in which the flies cannot distinguish between food containing 0% Zn and that containing 100% Zn and so consume food randomly from both diets, which limits egg production to the same level as when they only ingest food with 50% Zn.

At this same time point (day 8), *wDah* flies in the choice situation (0_100) also produced more eggs than the flies in the 0_0 condition (Figure 5B). However, unlike *rDah*, the flies with the 0_100 choice produced eggs at an equally high rate as flies on 100_100 (Figure 5B, Table S14). Surprisingly, *wDah* flies maintained on the 50_50 control diet pair also had higher than expected egg production, such that it was also indistinguishable from the positive control (100_100) (Figure 5B, Table S14). This indicates that dropping Zn to 50% of the level in the full feeding condition did not limit egg production of *wDah* females like it did for *rDah* females.

When assaying the flies on day 15, both genotypes showed the same egg-laying trends across food types as what they showed on day 8, but most differences between diet choice groups were reduced as egg laying dropped due to natural age-related decline in egg production (Figure S5 C-D, Table S14).

Together, these findings suggest that while dietary Zn levels can be physiologically limiting for egg production, the flies showed no evidence of selectively feeding on Zn-containing food to counteract the negative effects of Zn limitation on body stores or egg production.

Because of the way our assay is set up, we could also assess whether mothers used the presence of dietary Zn as a criterion for selecting the food on which they lay their eggs. When considering all our diet pairs, we found that the flies lay more eggs on Zn containing food than food lacking Zn (Figure 5C-D, Figure S5E-H, Table S15-S16). Although the strength of this egg site selection differed between genotypes, both demonstrated a stronger preference for Zn containing food the longer the assay continued. This analysis indicates that the flies express a diet-based choice for egg laying site selection that is sensitive to dietary Zn levels. However, it does not rule out the influence of other preferences that the flies may be expressing, which could produce the same outcome by chance. Examples include the tendency for conspecifics to lay eggs at the same site as a leading female (Sarin 2009, Moreira-Soto 2024) or some visible cues in the experimental set up that we are unaware of.

To test if the site selection was indeed specific to dietary Zn levels, we assessed if the flies’ laying site choices in the 0_100 condition significantly differed from egg laying site preferences demonstrated by flies with pairs of foods that were nutritionally identical. For both genotypes, there was little evidence that egg site selection for the flies in the 0_100 choice differed from egg laying site selection bias shown in controls after only 2 days of exposure to the choice (Figure S5E-F, Table S15-16). However, the strength of site selection in the 0_100 condition increased beyond any site bias in at least two of the three controls by day 8 (Figure 5C-D, Table S16) and of both controls by day 15 (Figure S5G-H, Table S16). Thus, both genotypes exhibited a growing strength of preference to lay eggs on food containing Zn over food lacking Zn over time. Given that maternal survival and feeding behaviour appear to be unaffected by dietary Zn depletion, this egg site preference may indicate that when maternal Zn stores drop, they select an egg laying site that protects larval survival, which relies on dietary Zn (Consuegra 2020).

### The *white* gene is required to maintain egg quality control during dietary Zn limitation

All of our data above show that the mutation in the *white* gene modifies egg laying and lifespan responses of flies to dietary Zn levels, albeit in relatively minor ways. One of the stronger phenotypes was the combined observation that in the choice assay, *wDah* females on food with 50% Zn laid eggs at the same rate as when on 100% Zn (Figure 5B), but they also retained a relatively high proportion of Zn in their body tissue when feeding on Zn depleted food (Figure 4B). We, therefore, wondered if *wDah* females had dysregulated egg production, such that they laid more, but lower quality, eggs when Zn was limited. To assess this, we measured the egg-to-adult viability of a sample of eggs collected on day 8 from all diet pairs used in the diet choice assay above (except 0_0 which produced insufficient eggs for the assay) (Figure 6).

**Figure 6.**
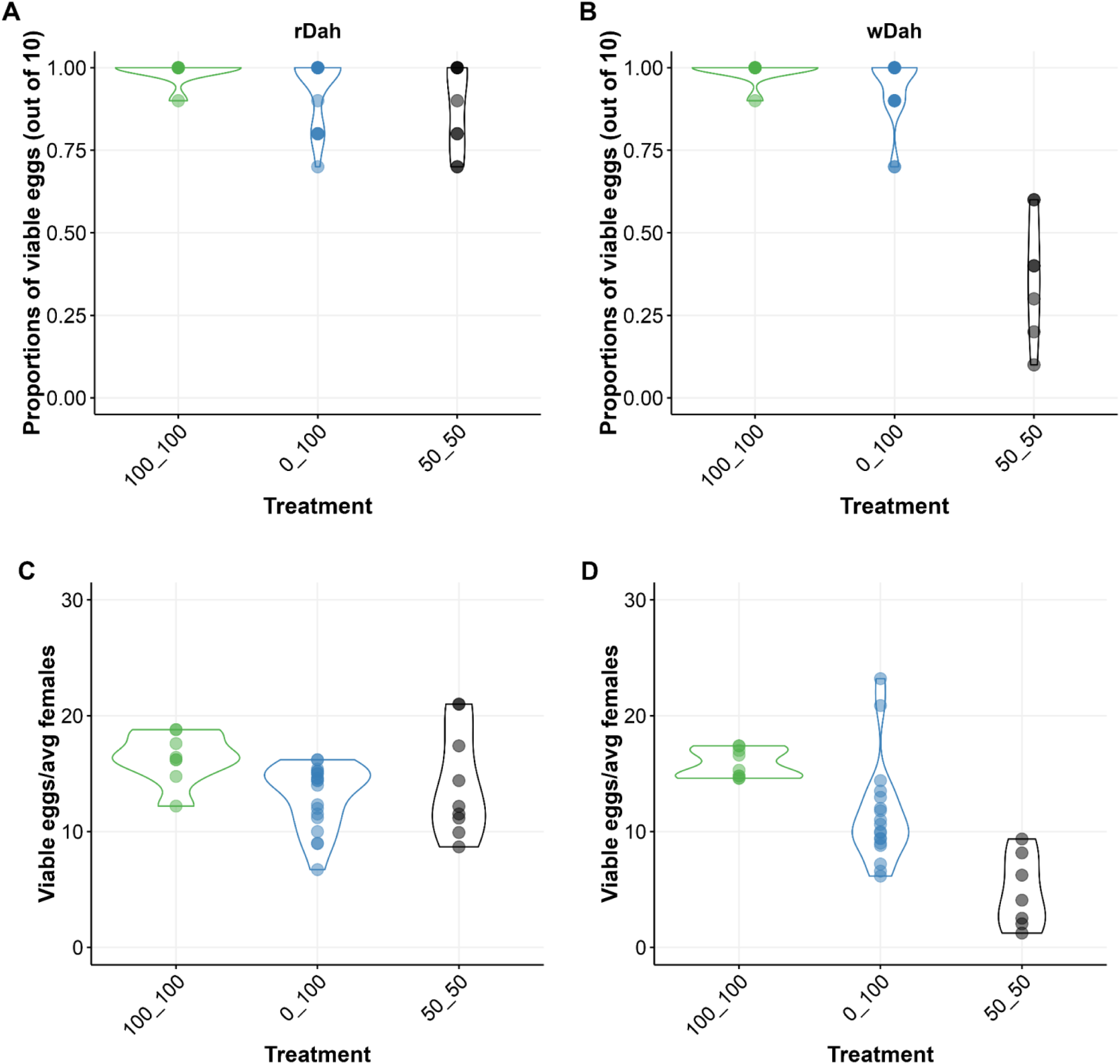
Egg viability is compromised when wDah, but not rDah, flies feed on Zn limiting diets. The proportion of viable eggs laid by (A) rDah mothers remains unchanged and high for flies that fed on any of the diet pairs tested in which Zn was manipulated (100_100, 0_100 or 50_50). By contrast, the proportion of viable eggs produced by (B) wDah mothers was significantly reduced by Zn limitation, which meant that, unlike (C) rDah mothers, (D) wDah females laid overall fewer viable eggs when feeding on Zn limiting diets.

Genotype significantly modified the proportion of viable eggs laid by females across the different diets (Figure 6A-B, Table S17). Specifically, while egg viability was unchanged and fixed at >80% for both genotypes maintained on most diet pairs, when *wDah* flies fed on the 50_ 50 diet, egg viability was significantly lower, reduced to ∼40% (Figure 6B, Table S18). This is the same condition in which *wDah* laid more eggs than anticipated (Figure 5B), meaning that while *rDah* flies produced the same absolute number of viable eggs in each food condition (Figure 6C, Table S19), *wDah* flies produced significantly fewer viable eggs when maintained under Zn limiting conditions (0_100 and 50_50) (Figure 6D, Table S19). Thus, although *wDah* mothers appear to benefit from the same regulatory mechanisms that protect maternal lifespan during dietary Zn depletion, the *white* gene appears to play an important role in their quality control system that maintains high quality egg production when dietary Zn levels fall.

## Discussion

Organisms require varying quantities and proportions of nutrients to grow, reproduce, and survive. The macronutrients are well characterised for playing a key role in determining fitness, especially for females who bear the larger anabolic load for offspring production (Lee 2008, Maklakov 2008, Boggs 2009, Simpson 2012). The micronutrients, which include vitamins and metal ions, are also critical for these processes, but their effects on reproduction and maternal physiology are less well characterised. Here, we explore the requirement for individual metal ions in *Drosophila* females and find that female flies employ different strategies in response to encountering food that is individually depleted for each of the different metal ions. Of these different responses, we focused on dietary Zn depletion because it elicited a unique combination of phenotypes, causing arrested egg production while preserving maternal lifespan. These data are consistent with dietary Zn being an important indicator of environmental quality that determines resource allocation between life histories that shape evolutionary fitness.

Life History Theory proposes that organisms partition limiting resources between reproduction and lifespan to optimize evolutionary fitness (Holliday 1989, Kirkwood 2000, Flatt 2018). Initially, these “resources” were assumed to be derived from the total amount of food, or energy, an organism eats, but more recently, it has become apparent that the relative abundance of specific nutrients in the diet determine the extent to which traits are expressed, independently of how much energy is in the diet (Simpson 2012). In particular, the protein : carbohydrate content of the diet has been repeatedly shown to be an important determinant of the inverse relationship between lifespan and reproduction (Lee 2008, Maklakov 2008, Skorupa 2008, Piper 2011, Fanson 2012, Solon-Biet 2014). Recently, we have characterised the role of a micronutrient in shaping these traits by studying how varying dietary sterols affect *Drosophila* females (Zanco 2021, Zanco 2022). Interestingly, even when mothers consume insufficient sterols to support sustained reproduction, they continue to take cues from dietary protein : carbohydrate proportions to determine egg production. Thus, to lay viable eggs, mothers must commit their own sterol reserves to those eggs, even if it shortens their lifespan. This finding is important because it shows that an essential dietary micronutrient could mediate the way that the macronutrients alter life histories. Here, our data show that another essential micronutrient, Zn, plays a key role in determining the way that egg production and lifespan are prioritised: Zn limitation preserves maternal lifespan and can even extend it in some of our trials and other recent work (Kosakamoto 2024), while causing reproduction to cease. Thus, Zn appears to be as important as the dietary macronutrients (protein and carbohydrate) in indicating food quality to reproducing females. Future experiments to determine the effect of the three-way interaction between protein, carbohydrate and Zn on lifespan and reproduction will be important to indicate the likely mechanistic underpinnings of Zn’s role in these phenotypes.

Given the importance of Zn in dictating reproduction, we were surprised to find that even after two weeks of Zn depletion, females did not alter their food choice behaviour to maximise egg output. This is not because the flies lack molecular sensors for Zn: they detect its loss and adjust their physiology to survive. Instead, it appears that their Zn sensing mechanism does not inform food choice decisions. This is especially surprising because we found that mothers prefer laying their eggs on food containing Zn over food without Zn. This is consistent with a strategy to ensure the success of their offspring. While *Drosophila* are known to exhibit nutrient-specific preferences that influence both feeding behaviour and oviposition (Edgecomb 1994, Lin 2019), the specific scenario where Zn deficiency impacts egg-laying substrate preference without altering food choice suggests distinct underlying mechanisms. Typically, sensory inputs such as olfactory and gustatory cues drive egg-laying site selection to optimize offspring survival, while the adult fly’s nutritional needs influence food choice (Yang 2008, Ribeiro 2010, Fujii 2015, Joseph 2015). The separation of these behaviours in the context of Zn sensing could be due to differences in the sensitivity of sensory receptors to Zn in food versus egg-laying substrates, or distinct neural processing pathways prioritizing reproductive success over immediate dietary intake. A recent study (Zhu 2015) investigated the neural mechanisms underlying egg-laying site selection in *Drosophila*, but it did not directly address Zn sensing. However, it provides evidence for the idea that distinct sensory pathways and circuits may be responsible for the uncoupling of Zn-dependent egg-laying preference and food choice. Future work to determine the neural circuits mediating these behaviours should reveal how this has been encoded.

One of the mechanistic findings in our data is that the *white* gene plays an important role in the way that dietary Zn restriction impacts on egg viability. *white* encodes a member of the ATP-binding cassette (ABC) transporter family, known for their role in transporting a wide array of substrates across cellular membranes (Mackenzie 1999, Dean 2001). Recent studies have highlighted the role of *white* in the Malpighian tubules, the fly kidneys, which is key for sequestering excess Zn ions, thus enabling Zn storage in the form of granules that can be retrieved for essential cellular processes as well as acting as a reservoir to prevent cytotoxicity if Zn is in excess (Mackenzie 1999, Yin 2017, Tejeda-Guzmán 2018). Mutants in the *white* gene exhibit reduced levels of stored Zn (Tejeda-Guzmán 2018). The specific mechanism by which the *white* gene influences this is not fully understood, but given its role in transmembrane transport, and its expression in the Malpighian tubules (Yin 2017), it may be involved in the active transport of Zn ions or Zn-bound complexes across the cellular membranes of the Malpighian tubules. The strongest phenotype we observed in our *white* mutated flies was a large reduction in the viability of eggs, but not their number, when dietary Zn was diluted to 50% of the level in our nutritionally complete food. This suggests that depleted Zn stores may lead to suboptimal Zn levels in the ovaries, which affects egg quality and viability, but not their production. To explore this hypothesis, conducting tissue-specific RNAi to knock down *white* expression selectively in the Malpighian tubules and measuring its effects on egg viability and Zn levels will be important.

Zn levels are particularly high in the *Drosophila* oocyte; a notable surge in Zn concentration acts as a signal for initiating oocyte maturation, which is crucial for successful fertilization and embryo growth (Hu 2020). When we deprived females of dietary Zn, or they are fed a diet containing the Zn-specific chelator TPEN, female egg production is reduced, presumably to avoid producing and laying inviable eggs (Hu 2020). Similar disruptions in oogenesis and decreased fertility have been observed in *C. elegans* hermaphrodites under Zn-restricted conditions (Hester 2017), and in humans, Zn deficiency has been linked to a range of adverse reproductive outcomes, including increased risks of infertility, miscarriage, and preterm delivery (Shah 2001, Nossier 2015). Zn deficiency is thought to be prevalent in almost all low- and middle-income countries, in part due to low environmental levels that give rise to Zn-poor crops (Gupta 2020). Thus, Zn limitation, and its effects on reproduction, may be a broadly relevant dietary selection pressure that flies have evolved to monitor in order to optimize their reproductive strategies.

## Conclusion

Our study demonstrates how dietary metal ions can affect both lifespan and reproduction in *Drosophila* females. In particular, we showed that the availability of dietary Zn determines the allocation of resources between reproduction and somatic maintenance, such that lifespan is preserved, even when dietary Zn availability is insufficient. Finally, we found an important role for the *white* gene, an eye pigment transporter, in controlling the production of viable eggs during Zn limitation. These data highlight the importance of metal ions in determining fly life histories and indicate that flies have evolved to include dietary Zn levels in their strategies to maximise reproductive success. Since dietary Zn levels affect reproduction in other organisms, including humans, these data could be useful in providing a platform for understanding how Zn determines whole organism health.

## Methods

### Fly husbandry

All experiments were conducted using two outbred “wild-type” *Drosophila* strains called Dahomey (abbreviated here as *rDah*) and white Dahomey (*wDah*). These strains have the same genetic background, but the latter is a homozygous mutant for the *white* gene, which causes the flies to have white eyes (Bingham 1980, Hazelrigg 1984, Mair 2005). White-eyed flies mutant for this gene are commonly used as the genetic background for transgenesis since they provide an easy-to-visualise selectable marker to distinguish between transgenics that carry a construct to complement the mutation (orange to red-eyed) and controls (white-eyed). *wDah* and *rDah* stocks are maintained in large numbers in continuous overlapping generations in a high-density population cage at a constant temperature of 25°C, under 12-hr light: dark photoperiods. Upon removal from the population cages, flies were reared for two generations at a controlled density using the eggs laid by age-matched mothers before use in experiments, to control for possible parental effects (Linford 2013). Following the eclosion of the third generation, newly emerged adults were allowed to mate for 48 hours before they were lightly anaesthetised with CO_2_ and sorted by sex. All stocks were maintained on sugar yeast food (Bass 2007).

### Experimental diets

The completely defined, synthetic (holidic) diets were prepared using the exome matched FLYAA formula as described by *Piper et al.* (Piper 2014, Piper 2017) (Table S20). For results shown in Figure 1, four different experimental diets were created by diluting down the mixture of all metal ions to 0%, 10%, and 50% of the level in the complete (control) diet. Diets for Figure 2 were prepared by dropping each metal ion separately (Ca, Cu, Fe, Mg, Mn, and Zn). For Figure 4-6, Zn was diluted to four different concentrations (0%, 10%, 50% and 100%) of the original stock solution (Table S20-21).

### Lifespan assays

For each experimental diet, 10 female flies were placed into each of ten vials per genotype. Every 2-3 days, flies were transferred into new vials containing fresh food at which point deaths and censors were recorded (Piper 2016) and saved using the software Dlife (Linford 2013). 10 replicates of 10 flies per vial were used per treatment diet.

### Fecundity assays

For the fecundity assay, digital images of the surface of the food with eggs were acquired using a web camera mounted on a Zeiss dissecting microscope and eggs were counted manually from the images. Egg production was recorded on days 8 and 15 of the experiment after the flies had been exposed to the diets for 24 hours. Fecundity was measured as the number of eggs laid per female during each laying period. 10 replicates of 10 flies per vial were used per treatment diet.

### Quantification of metal ions

To assess the impact of Zn deprivation on metal ion levels, we separated ovaries from the remaining body tissue of *rDah* and *wDah* flies after 8 days of feeding on Zn-deficient and complete (100N) diet. On the 8th day of adulthood, the flies were anesthetized using CO₂, and the ovaries were carefully dissected from the remaining body tissue. Both ovaries and body tissues were collected separately for each genotype. The collected samples were freeze-dried for a few days until completely dry. Each dried sample was then treated with 50 µL of 65% HNO₃ and left overnight at room temperature. The samples were subsequently heated at 90°C for 20 minutes, followed by the addition of 50 µL of H₂O₂ and further heating at 70°C for 15 minutes. The digested samples were diluted to a final volume of 1 mL with deionized water. Metal ion levels (Zn, Cu, Fe, Mg, and Mn) were measured using an Agilent 8800 Triple Quad Inductively Coupled Plasma-Mass Spectrometer (ICP MS). Calibration was performed using standard solutions for each metal (Figure 4). 5 replicates of ovary and body tissues were used for the two treatment diets.

### Zinc Choice assay

Flies for the Zn choice assay were generated in the same way as for the lifespan assay. Treatment conditions consisted of two *Drosophila* maintenance vials, each containing food at their base, with their openings taped together with electrical tape so that flies could freely walk between the two ends to choose their food (Figure S4). The food pairs were a choice condition containing 0% Zn at one end of the vial pair, and 100% Zn at the other (0_100); a positive control with 100% Zn food (complete diet) (100_100) at both ends; a negative control with 0% Zn food at both ends (0_0) and; a control for random sampling with 50% Zn at both ends (50_50). To control for side preferences unrelated to food composition, vial orientation was noted for each vial pair. Five flies per genotype were placed into each of the ten connected vial pairs for each treatment condition. Egg number was counted in each vial on days 2, 8 and 15 of adulthood after a 24 hour egg-laying period and the side on which each egg was laid was noted.

### Egg-to-adult viability assay

Egg-to-adult viability from each of the choice assay groups was assessed by transferring 10 randomly selected eggs from the diet choice vials to vials containing SY food, which is optimal for fly development. The total number of adults that emerged was counted across the replicate vials for each of the treatments and divided by the number of eggs to give the proportion viable. We ensured equal distribution of eggs from both ends of the diet pairs into separate SY vials. In cases where there weren’t enough eggs from one end, we combined eggs from both ends of a diet pair into one SY vial.

### Statistical analyses

Statistical analyses were performed using the R Version 2023.06 across various experiments. For survival analyses, linear models were employed to analyze median lifespan. The significance of relationships between genotypes and treatments was evaluated using Type II ANOVA from the package car (Fox 2019). Post-hoc pairwise comparisons (Bonferroni adjusted) were conducted for genotypes to evaluate median lifespan differences.

Fecundity responses to metals, and Zn concentrations were analyzed using Linear Mixed-Effects Models (LMMs) with the lmer function (Bates 2015). These models assessed fixed effects for day, dilution level, genotype, and their interactions, alongside random effects for replicate variability. Fecundity responses to individual metal ions were also analyzed similar as above, including variables such as day, treatment, and genotype. Type III ANOVA (Fox 2019) was used to assess the significance of fixed effects and interactions.

Egg-to-adult viability on metal dropouts was analyzed using zero-inflated Poisson models fitted with the glmmTMB function (Brooks 2017). The models included fixed effects for metal dropout, genotype, and their interaction, with Type II ANOVA (Fox 2019) assessing the significance of these effects.

ICP MS analysis of metal concentrations employed linear models with Proportion as the response variable and metal, genotype, and tissue as predictors, including their interactions. Type III ANOVA (Fox 2019) evaluated the overall impact and interactions of these factors, with post-hoc pairwise comparisons adjusted using the Tukey method.

In the Zn selective feeding assay, linear mixed-effects models were used to analyze the impact of dietary Zn variations on egg production. These models incorporated fixed effects for treatment, genotype, day, and their interactions, alongside random effects and diet side for experimental blocks. The significance of fixed effects was evaluated using Type III ANOVA (Fox 2019), with post-hoc pairwise comparisons conducted using the emmeans package (Lenth 2021).

To assess whether flies prefer laying eggs in food that contains Zn over food that does not, we first fit the data from the choice scenario (0_100) with a generalized linear mixed effects model (GLMMs) using the glmmTMB function (Brooks 2017) using a binomial distribution. In this model, we used the number of flies in the vial as a covariate, day, genotype and their interactions as fixed effects, and replicate vial and block as random effects. We tested the model for significant fit using a Type III ANOVA (Fox 2019), then tested if the distribution of eggs across the two vials was significantly different from no choice (assuming a null mean of 0.5) using post hoc tests from the emmeans package (Lenth 2021). This approach allowed us to determine if females show a preference for laying their eggs in Zn-containing food, and how this preference changes over time and differs between genotypes.

To be sure that our no choice scenario was not driven by the fact that females tend to lay their eggs together with other eggs, we then compared the choice scenario (0_100) against three no choice controls (0_0, 50_50, and 100_100). For each of the control diets, we randomly assigned one of the two vials from each replicate to the treatment group using a custom built script in R (see R scripts in FigShare - DOI: 10.26180/26550244). To do this, we employed a generalized linear mixed effects model (GLMMs) using the glmmTMB function (Brooks 2017) with a binomial distribution. This model included the total number of flies in a vial as a covariate, day, diet, genotype and their interactions as fixed effects, and replicate vial and block as random effects. After testing the model for a significant fit using a Type III ANOVA (Fox 2019), we compared the choice treatment (0_100) to the three controls (0_0, 50_50, and 100_100) using trt.vs.ctrl custom contrasts in the emmeans package (Lenth 2021). This approach allowed us to determine if the choice treatment resulted in a different distribution of egg laying sites than the three control treatments.

For the analysis of egg-to-adult viability across different dietary Zn combinations and genotypes, a generalized linear mixed model (GLMM) using the glmmTMB function (Brooks 2017) was employed. The model was specified with a binomial family and logit link to handle the binary nature of the viability data, which consisted of counts of emerged and non-emerged eggs. The model included fixed effects for treatment and genotype, and random effects. The significance of the fixed effects was evaluated using an ANOVA Type III test via the Anova function (Fox 2019). Post-hoc comparisons were conducted using the emmeans function, followed by pairwise comparisons.

For the analysis of viable eggs per average female, a zero-inflated Poisson model was utilized, implemented using the glmmTMB function (Brooks 2017). The significance of the fixed effects was evaluated using a Type III ANOVA (Fox 2019), and subsequent post-hoc analyses were conducted to explore specific differences between treatments and genotypes.

Plots were produced using ggplot2 (Wickham 2016).

## Authors’ contributions

S.S: Conceptualization, data curation, formal analysis, investigation, methodology, visualization, writing—original draft. H.T.H.T: Data curation, investigation. G.M: Data curation, investigation. R.B: Conceptualization, supervision, writing—review & editing. C.K.M: Conceptualization, formal analysis, funding acquisition, methodology, supervision and writing—review and editing. M.D.W.P: Conceptualization, funding acquisition, methodology, resources, supervision and writing—review and editing.

## Acknowledgements

We thank Mia Wansborough, Tahlia Fulton, Joshua Johnstone and Jiayi Lin (Monash University) for their invaluable assistance in sorting flies for lifespan studies. We thank Tahlia Fulton (Monash University) for helping with some of the preliminary data analysis. Additionally, we thank Fumiaki Obata and Hina Kosakamoto (RIKEN Center for Biosystems Dynamics Research) for their discussions related to this work.

