## Supplemental table for "Dietary zinc limitation dictates lifespan and reproduction trade-offs of *Drosophila* mothers": Supplemental tables.docx

Table S1.
Anova (Type II) on linear model to best represent the effects of genotypes and dietary treatments (metal dilution diets represented as metal concentration) on median lifespan.

|  | Sum Sq | Df | *P value* |
| --- | --- | --- | --- |
| Genotype | 357 | 1 | 0.181 |
| Treatment | 367328 | 3 | <0.001^***^ |
| Genotype: Treatment | 5292 | 3 | <0.001^***^ |

Table S2.

Post-hoc pairwise (Bonferroni adjusted) comparisons of genotypes to evaluate median lifespan differences between 100% metal diet (control) and 50%, 10% and 0% metal diets respectively

Genotype= *rDah*

| Contrast | Estimate | df | *p value* |
| --- | --- | --- | --- |
| 100% - 50% | -0.461 | 714 | 0.9963 |
| 100% - 10% | 43.523 | 714 | <0.001^***^ |
| 100%-0% | 55.353 | 714 | <0.001^***^ |

Genotype= *wDah*

| Contrast | Estimate | df | *p value* |
| --- | --- | --- | --- |
| 100% - 50% | -2.741 | 714 | 0.5575 |
| 100% - 10% | 30.639 | 714 | <0.001^***^ |
| 100% - 0% | 44.688 | 714 | <0.001^***^ |

Table S3.
Anova (type III) of the linear model to best represent the relationship between the days on which eggs (age) were counted, genotypes and dietary metal ion concentration

|  | Chisq | Df | *P value* |
| --- | --- | --- | --- |
| Age | 50.92 | 1 | <0.001^***^ |
| Metal Concentration | 1418.05 | 3 | <0.001^***^ |
| Genotype | 3.52 | 1 | 0.06 |
| Age: Metal Concentration | 20.73 | 3 | <0.001^***^ |
| Age: Genotype | 6.30 | 1 | <0.05^*^ |
| Metal Concentration: Genotype | 32.49 | 3 | <0.001^***^ |
| Age: Metal Concentration: Genotype | 7.87 | 3 | <0.05^*^ |

Table S4.

Anova (Type II) on linear model to best represent the relationship between genotypes, treatment (metal dropout diets) and median lifespan

|  | Sum Sq | Df | *P value* |
| --- | --- | --- | --- |
| Genotype | 21994 | 1 | <0.001^***^ |
| Treatment | 158452 | 6 | <0.001^***^ |
| Genotype: Treatment | 8090 | 6 | <0.001^***^ |

Table S5.

Post-hoc pairwise comparisons (Bonferroni adjusted) for genotypes to evaluate median lifespan differences between All metal diet (control) and different metal dropout diets

Genotype= *rDah*

| Contrast | Estimate | df | *p value* |
| --- | --- | --- | --- |
| All metals - (-Cu) | 1.055 | 1275 | 1.0000 |
| All metals - (-Ca) | 5.938 | 1275 | 0.2793 |
| All metals - (-Fe) | 3.934 | 1275 | 1.0000 |
| All metals - (-Mg) | 35.501 | 1275 | <0.001^***^ |
| All metals - (-Zn) | -4.258 | 1275 | 1.0000 |
| All metals - (-Mn) | 5.154 | 1275 | 0.7158 |

Genotype= *wDah*

| Contrast | Estimate | df | *p value* |
| --- | --- | --- | --- |
| All metals - (-Cu) | 6.256 | 1275 | 0.2841 |
| All metals - (-Ca) | 5.499 | 1275 | 0.5159 |
| All metals - (-Fe) | 9.810 | 1275 | <0.01^**^ |
| All metals - (-Mg) | 26.207 | 1275 | <0.0001^***^ |
| All metals - (-Zn) | -8.513 | 1275 | <.05^*^ |
| All metals - (-Mn) | 2.461 | 1275 | 1.0000 |

Table S6.
Anova (type III) of the linear model to best represent the relationship between the days of adulthood (age) when eggs were counted, genotypes and treatment (metal dropout diets) and their interactions. Higher-order interaction terms were removed in a stepwise fashion when removal did not significantly affect model fit (determined by AIC).

|  | Chisq | Df | *P value* |
| --- | --- | --- | --- |
| Age | 471.196 | 1 | <0.001^***^ |
| Treatment | 2468.271 | 6 | <0.001^***^ |
| Genotype | 17.559 | 1 | <0.001^***^ |
| Age: Treatment | 167.327 | 6 | <0.001^***^ |
| Age: Genotype | 9.185 | 1 | <0.01^**^ |

Table S7.

Anova (type II) of the zero-inflated model to best represent the relationship between egg-to-adult viability of eggs collected from different metal diets and genotypes.

|  | Chisq | Df | *P value* |
| --- | --- | --- | --- |
| Metal dropout | 5.83 | 4 | 0.2119 |
| Genotype | 1.33 | 1 | 0.2496 |
| Metal dropout: Genotype | 3.57 | 4 | 0.4668 |

Table S8.
Anova (Type II) on linear model to best represent the relationship between genotypes, treatment (zinc dilution diets represented as zinc concentration) and median lifespan

|  | Sum Sq | Df | *p value* |
| --- | --- | --- | --- |
| Genotype | 2752 | 1 | 0.004^**^ |
| Treatment | 1606 | 3 | 0.184 |
| Genotype: Treatment | 1119 | 3 | 0.338 |

Table S9.

Post-hoc pairwise comparisons (Bonferroni adjusted) for genotypes to evaluate lifespan differences between 100% zinc diet (control) and 50%, 10% and 0% zinc diets respectively

Genotype= *rDah*

| Contrast | Estimate | df | *p value* |
| --- | --- | --- | --- |
| 100% - 50% | 2.957 | 678 | 1.0000 |
| 100% - 10% | -2.807 | 678 | 1.0000 |
| 100% - 0% | 0.825 | 678 | 1.0000 |

Genotype= *wDah*

| Contrast | Estimate | df | *p value* |
| --- | --- | --- | --- |
| 100% - 50% | -2.196 | 678 | 1.0000 |
| 100% - 10% | -2.816 | 678 | 1.0000 |
| 100% - 0% | 2.268 | 678 | 1.0000 |

Table S10.
Anova (type III) of the linear model to best represent the relationship between the days of adulthood (age) the eggs counted, genotypes and dietary zinc concentration.

|  | Chisq | Df | *p value* |
| --- | --- | --- | --- |
| Age | 235.34 | 1 | <0.001^***^ |
| Zinc Concentration | 825.10 | 3 | <0.001^***^ |
| Genotype | 24.07 | 1 | <0.001^***^ |
| Age: Zinc Concentration | 56.00 | 3 | <0.001^***^ |
| Age: Genotype | 0.16 | 1 | 0.68 |
| Zinc Concentration: Genotype | 15.65 | 3 | <0.01^**^ |
| Age: Zinc Concentration: Genotype | 16.67 | 3 | <0.001^***^ |

Table S11.

Anova (type III) of the linear model representing the relationship between the proportions of metals measured (where each metal is expressed as a proportion of that flies fed on 0Zn diets, divided by that found in flies feeding on a complete diet), tissues (body & ovary) and genotypes (*rDah* and *wDah*)

|  | Sum Sq | Df | *p value* |
| --- | --- | --- | --- |
| Metal | 16.166 | 4 | <0.001^***^ |
| Genotype | 0.211 | 1 | <0.001^***^ |
| Tissue | 10.566 | 1 | <0.001^***^ |
| Metal: Genotype | 1.596 | 4 | <0.001^***^ |
| Metal: Tissue | 5.688 | 4 | <0.001^***^ |
| Genotype: Tissue | 0.196 | 1 | <0.001^***^ |
| Metal: Genotype: Tissue | 0.522 | 4 | <0.001^***^ |

Table S12.

Differences in metal levels across tissues and genotypes, measured using Tukey’s multiple comparisons tests

| Tissue | Metal | Df | emmean | Group |
| --- | --- | --- | --- | --- |
| *Body* | *rDah* | | | |
|  | Zn | 480 | 0.153 | 1 |
|  | Fe | 480 | 0.917 | 3 |
|  | Mg | 480 | 1.029 | 4 5 6 |
|  | Mn | 480 | 1.069 | 5 6 |
|  | Cu | 480 | 1.126 | 6 |
|  | *wDah* | | | |
|  | Zn | 480 | 0.528 | 2 |
|  | Fe | 480 | 0.993 | 3 4 5 |
|  | Mg | 480 | 1.069 | 5 6 |
|  | Mn | 480 | 0.954 | 3 4 |
|  | Cu | 480 | 1.256 | 7 |
| *Ovary* | *rDah* | | | |
|  | Zn | 480 | 0.362 | 1 |
|  | Fe | 480 | 0.223 | 4 |
|  | Mg | 480 | 0.203 | 3 4 |
|  | Mn | 480 | 0.145 | 2 3 4 |
|  | Cu | 480 | 1.126 | 3 4 |
|  | *wDah* | | | |
|  | Zn | 480 | 0.101 | 1 2 |
|  | Fe | 480 | 0.132 | 1 2 3 4 |
|  | Mg | 480 | 0.156 | 2 3 4 |
|  | Mn | 480 | 0.112 | 1 2 3 |
|  | Cu | 480 | 1.256 | 2 3 4 |

Table S13.

Anova (type III) of the linear mixed model to best represent the relationship between egg production, treatment (diet combinations with dietary zinc variations), days of adulthood (age) and genotype.

|  | Chisq | Df | *p value* |
| --- | --- | --- | --- |
| Treatment | 196.90 | 3 | <0.001^***^ |
| Genotype | 17.12 | 1 | <0.001^***^ |
| Age | 1221.46 | 2 | <0.001^***^ |
| Treatment: Genotype | 10.32 | 3 | 0.016^*^ |
| Treatment: Age | 81.02 | 6 | <0.001^***^ |
| Genotype: Age | 40.29 | 2 | <0.001^***^ |
| Treatment: Genotype: Age | 6.50 | 6 | 0.369 |

Table S14.

Differences between egg production on different treatments (diet combinations with dietary zinc variations) counted on different days of adulthood (age) and genotypes measured using Tukey’s multiple comparisons tests

| Age | Treatment | emmean | Group |
| --- | --- | --- | --- |
| 2 | *rDah* | | |
|  | 0_100 | 32.657 | 1 |
|  | 0_0 | 32.838 | 1 |
|  | 50_50 | 34.692 | 12 |
|  | 100_100 | 39.492 | 2 |
|  | *wDah* | | |
|  | 0_100 | 27.560 | 1 |
|  | 0_0 | 24.598 | 1 |
|  | 50_50 | 27.998 | 1 |
|  | 100_100 | 29.548 | 1 |
| 8 | *rDah* | | |
|  | 0_100 | 17.503 | 2 |
|  | 0_0 | 2.968 | 1 |
|  | 50_50 | 19.959 | 2 |
|  | 100_100 | 28.128 | 3 |
|  | *wDah* | | |
|  | 0_100 | 17.503 | 2 |
|  | 0_0 | 2.968 | 1 |
|  | 50_50 | 19.959 | 3 |
|  | 100_100 | 28.128 | 23 |
| 15 | *rDah* | | |
|  | 0_100 | 6.683 | 2 |
|  | 0_0 | 0.615 | 1 |
|  | 50_50 | 6.065 | 12 |
|  | 100_100 | 9.278 | 2 |
|  | *wDah* | | |
|  | 0_100 | 8.867 | 2 |
|  | 0_0 | 1.192 | 1 |
|  | 50_50 | 9.288 | 2 |
|  | 100_100 | 10.105 | 2 |

Table S15.

Analysis of Deviance (Type III Wald Chisquare Tests) of the generalized linear mixed model to represent the relationship between egg-laying preferences and factors including total flies, treatment conditions, days of adulthood (Age), and genotypes

|  | Chisq | Df | *p value* |
| --- | --- | --- | --- |
| Total_fly | 5.34 | 1 | <0.05^*^ |
| Treatment | 4.45 | 3 | 0.217 |
| Age | 13.61 | 2 | <0.01^**^ |
| Genotype | 25.59 | 1 | <0.001^***^ |
| Treatment: Day | 214.19 | 6 | <0.001^***^ |
| Treatment: Genotype | 122.07 | 3 | <0.001^***^ |
| Age: Genotype | 1.59 | 2 | 0.451 |
| Treatment: Day: Genotype | 83.66 | 6 | <0.001^***^ |

Table S16.

Comparing egg-laying preferences across diet combinations with dietary Zn Variations, days of adulthood (Age), and genotypes using Tukey's multiple comparisons tests

| Age | Contrast | Estimate | df | *p value* |
| --- | --- | --- | --- | --- |
| 2 | *rDah* | | | |
|  | (0_0)- (0_100) | -0.057 | Inf | 0.976 |
|  | (100_100)- (0_100) | -0.461 | Inf | 0.115 |
|  | (50_50)- (0_100) | -0.038 | Inf | 0.989 |
|  | *wDah* | | | |
|  | (0_0)- (0_100) | -0.042 | Inf | 0.9874 |
|  | (100_100)- (0_100) | -0.969 | Inf | <0.001^***^ |
|  | (50_50)- (0_100) | 0.429 | Inf | 0.164 |
| 8 | *rDah* | | | |
|  | (0_0)- (0_100) | -0.707 | Inf | <0.05^*^ |
|  | (100_100)- (0_100) | -0.239 | Inf | 0.598 |
|  | (50_50)- (0_100) | -0.894 | Inf | <0.001^***^ |
|  | *wDah* | | | |
|  | (0_0)- (0_100) | -0.579 | Inf | 0.102 |
|  | (100_100)- (0_100) | -1.453 | Inf | <0.001^***^ |
|  | (50_50)- (0_100) | -1.501 | Inf | <0.001^***^ |
| 15 | *rDah* | | | |
|  | (0_0)- (0_100) | -1.497 | Inf | <0.001^***^ |
|  | (100_100)- (0_100) | -0.594 | Inf | <0.05^*^ |
|  | (50_50)- (0_100) | -1.179 | Inf | <0.001^***^ |
|  | *wDah* | | | |
|  | (0_0)- (0_100) | -1.767 | Inf | <0.001^***^ |
|  | (100_100)- (0_100) | -1.094 | Inf | <0.001^***^ |
|  | (50_50)- (0_100) | -1.067 | Inf | <0.001^***^ |

Table S17.

Anova (type III) of the zero-inflated model to best represent the relationship between egg-to-adult viability of eggs collected from different treatments (diet combinations with dietary Zn variations), and genotypes.

|  | Chisq | Df | *p value* |
| --- | --- | --- | --- |
| Treatment | 5.06 | 2 | 0.079 |
| Genotype | 0.01 | 1 | 0.903 |
| Treatment: Genotype | 17.14 | 2 | <0.001^***^ |

Table S18.

Differences in the proportion of eggs that gave rise to viable adults across different treatments (diet combinations with dietary Zn variations). Data analyses were performed on genotypes separately using Tukey’s multiple comparisons tests.

Genotype= *rDah*

| Treatment | emmean | Group |
| --- | --- | --- |
| 100_100 | 4.655 | 1 |
| 0_100 | 3.017 | 1 |
| 50_50 | 2.154 | 1 |

Genotype= *wDah*

| Treatment | emmean | Group |
| --- | --- | --- |
| 100_100 | 4.783 | 2 |
| 0_100 | 2.960 | 2 |
| 50_50 | -0.485 | 1 |

Table S19.

Differences in the absolute number of viability eggs across different treatments (diet combinations with dietary Zn variations), and genotypes measured using Tukey’s multiple comparisons tests

Genotype= *rDah*

| Treatment | emmean | Group |
| --- | --- | --- |
| 100_100 | 2.77 | 1 |
| 0_100 | 2.56 | 1 |
| 50_50 | 2.58 | 1 |

Genotype= *wDah*

| Treatment | emmean | Group |
| --- | --- | --- |
| 100_100 | 2.76 | 3 |
| 0_100 | 2.38 | 2 |
| 50_50 | 1.40 | 1 |

Table S20.

Stock Solution for metal ions

| FLYAA (Complete holidic diet- also referred to as 100N) | |
| --- | --- |
| Ingredients/Nutrients | Quantity/L |
| MilliQ Water | 777 ml |
| Agar | 7 g |
| L-isoleucine | 0.56 g |
| L-leucine | 1.02 g |
| L-tyrosine | 0.46 g |
| Sucrose | 17.12 g |
| Cholesterol | 15 ml |
| CaCl2 | 1 ml |
| CuSO4 | 1 ml |
| FeSO4 | 1 ml |
| MgSO4 | 1 ml |
| MnCl2 | 1 ml |
| ZnSO4 | 1 ml |
| Acetate Buffer | 100 ml |
| Nucleic Acid and lipids | 8 ml |
| Essential Amino Acids (EAA) | 30.2 ml |
| Non-Essential Amino Acids (NEAA) | 30.2 ml |
| Glutamate | 7.59 ml |
| Cysteine | 3.41 ml |
| Vitamin | 21 ml |
| Folic Acid | 1 ml |
| Nipagin | 15 ml |
| Propionic Acid | 6 ml |

| Metal ions in Holidic diet | g/l | Final Concentration (g/l) | Solubility (g/l) | Metal ion in solution (g/l) |
| --- | --- | --- | --- | --- |
| CaCl2.6H2O | 250 | 0.25 | 745 | 0.0680 |
| MgSO4 | 250 | 0.25 | 255 | 0.0498 |
| CuSO4.5H2O | 2.5 | 0.0025 | 316 | 0.0006 |
| FeSO4.7H2O | 25 | 0.025 | soluble | 0.0050 |
| MnCl2.4H2O | 1 | 0.001 | 723 | 0.0003 |
| Zn SO4.7H2O | 25 | 0.025 | 300 | 0.0057 |

Table S21.

Modified holidic diets (FLYAA)

| Diet | Modifications |
| --- | --- |
| Calcium dropout (-Ca) | Diet made without adding calcium |
| Magnesium dropout (-Mg) | Diet made without adding magnesium |
| Copper dropout (-Cu) | Diet made without adding copper |
| Iron dropout (-Fe) | Diet made without adding Iron |
| Manganese dropout (-Mn) | Diet made without adding Manganese |
| Zinc dropout (-Zn) | Diet made without adding Zinc |
| 100% metals/ 100% Zn/ All metals/ 100N | Complete diet. No manipulations |
| 50% metals | Each metal added to the diet were diluted to 50% of their original stock solution |
| 10% metals | Each metal added to the diet were diluted to 10% of their original stock solution |
| 0% metals | No metals were added to the diet |
| 50% Zn | The zinc added to the diet was diluted to 50% of their original stock solution |
| 10% Zn | The zinc added to the diet was diluted to 10% of their original stock solution |
| 0% Zn | No zinc was added to the diet |

^[1]^

1. For 10% dilution: Add 5ml of stock solution to 45ml of miliQ water to make a 50ml diluted solution.

   For 50% dilution: Add 25ml of stock solution to 25ml of miliQ water to make a 50ml diluted solution.

   For 100% dilution: Substitute the metal ions in the holidic food (e.g., zinc, iron, copper, manganese, and magnesium) with 1ml of miliQ water for each in 1L holidic food.

   For 0% dilution: Follow the 100% recipe without any changes.
