## Supplemental Figures for "Dietary zinc limitation dictates lifespan and reproduction trade-offs of *Drosophila* mothers": Supplemetal figures.docx

**Figure S1**

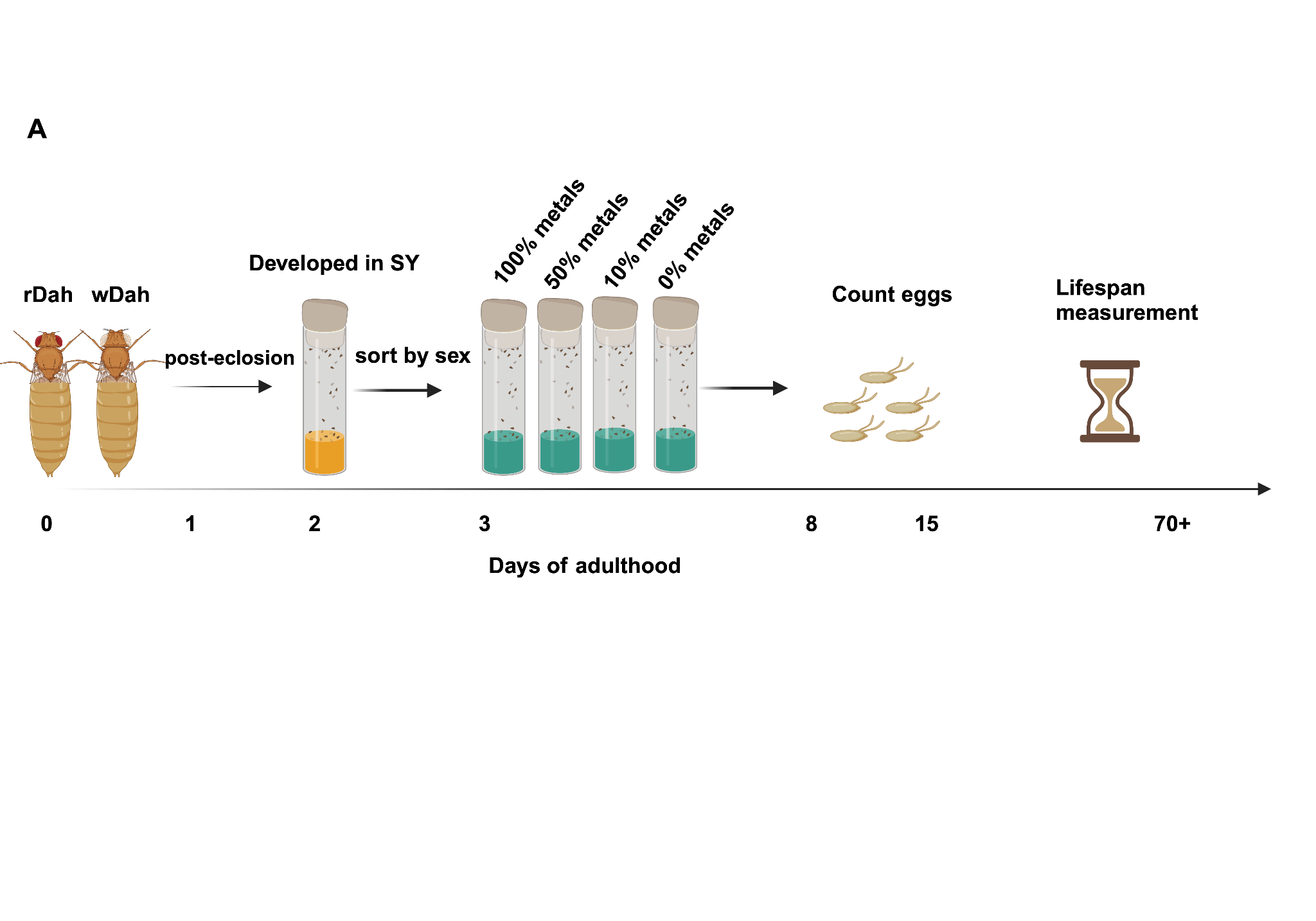

**Figure S1. Experimental timeline and dietary regimens for adult female flies for data shown in Figure 1**

*A) Flies of two genotypes, rDah and wDah were maintained separately on SY (sugar / yeast) food until thethird day of adulthood. Once mated, on day 3, they were each transferred to four different synthetic diets and reproduction and lifespan measured throughout life. Diets included a complete diet (100 % metals), 50% metals, 10% metals and 0% metals. N= 100 flies/diet*

**Figure S2**

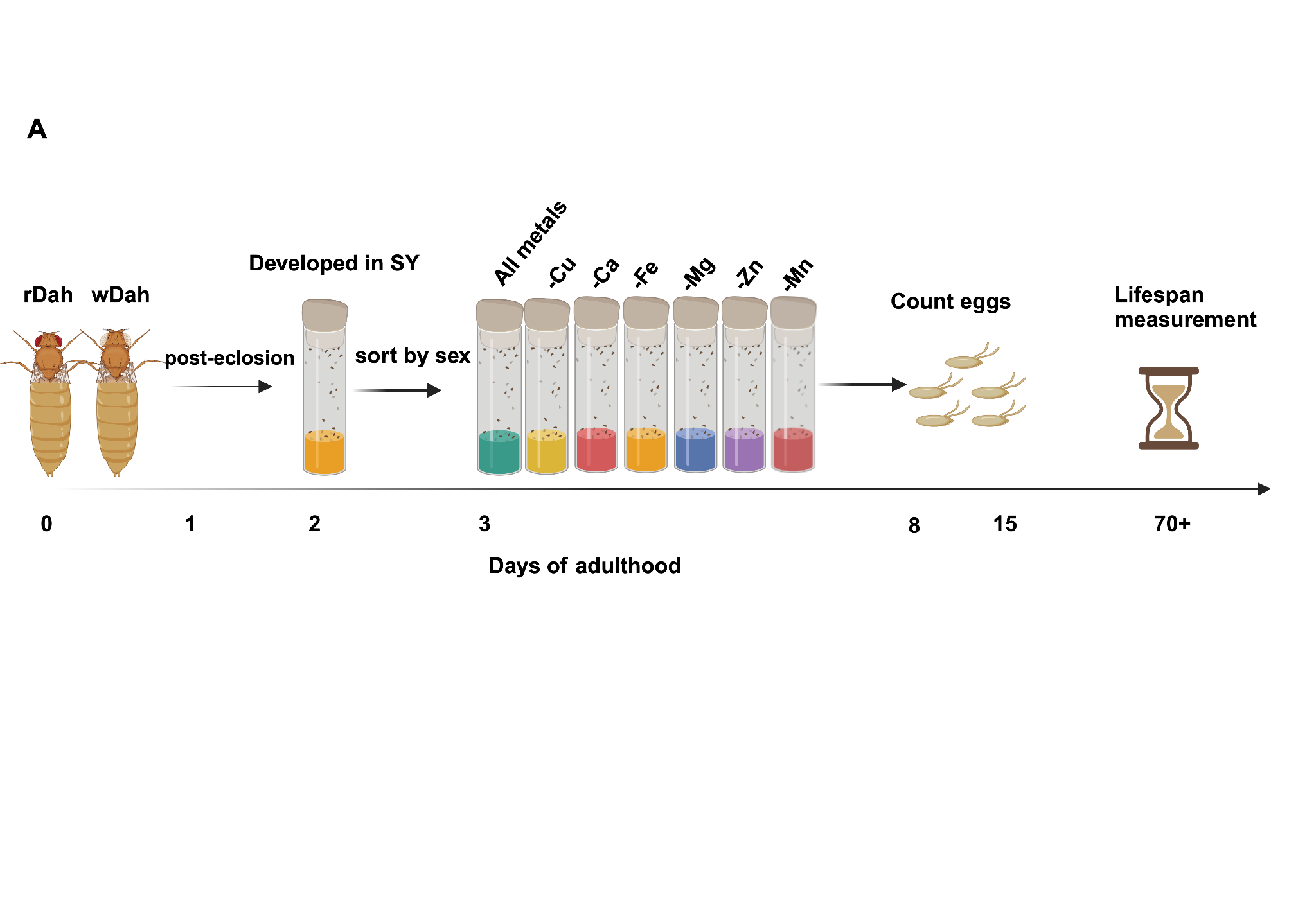

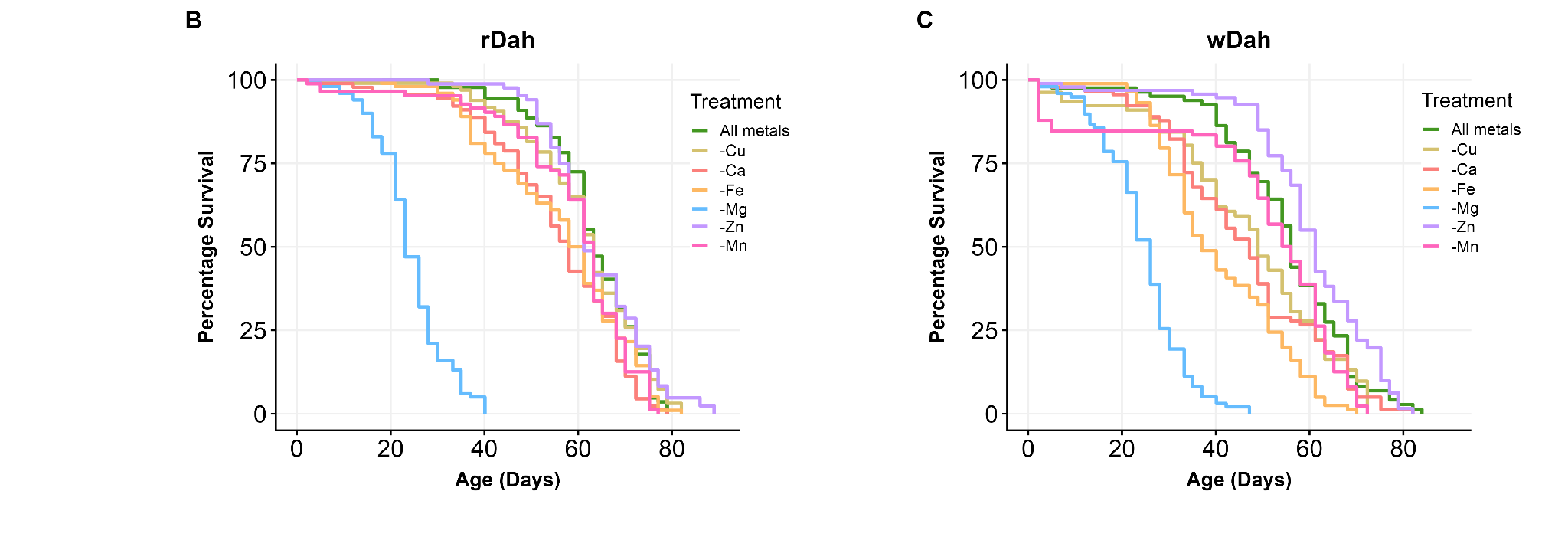

**Figure S2. Experimental timeline, dietary regimens, and lifespan analysis for adult female flies (data shown in Figure 2): Genotype-dependent responses to metal dropout diets**

*A) Flies of two genotypes, rDah and wDah were maintained separately on SY (Sugar / Yeast) food until the third day of adulthood. Once mated, on day 3, they were each transferred to seven different synthetic diets. Diets included a complete diet (all metals), a Ca, Cu, Fe, Mg, Mn, and Zn dropout diets. N= 100 flies/diet*

*B) The lifespan responses of rDah to individual metal depletion were minor, except for Mg deprivation, which resulted in severe lifespan shortening. No significant difference in lifespan was observed for flies deprived of Cu, Zn, Mn, and Fe.*

*C)The responses of wDah varied more, as Fe, Mg and Zn all significantly modified lifespan to some extent. A mild but significant lifespan extending effect when deprived of Zn compared to flies fed a complete diet. Lifespan shortening effects were observed when depleted of Mg, Cu, and Mn.*

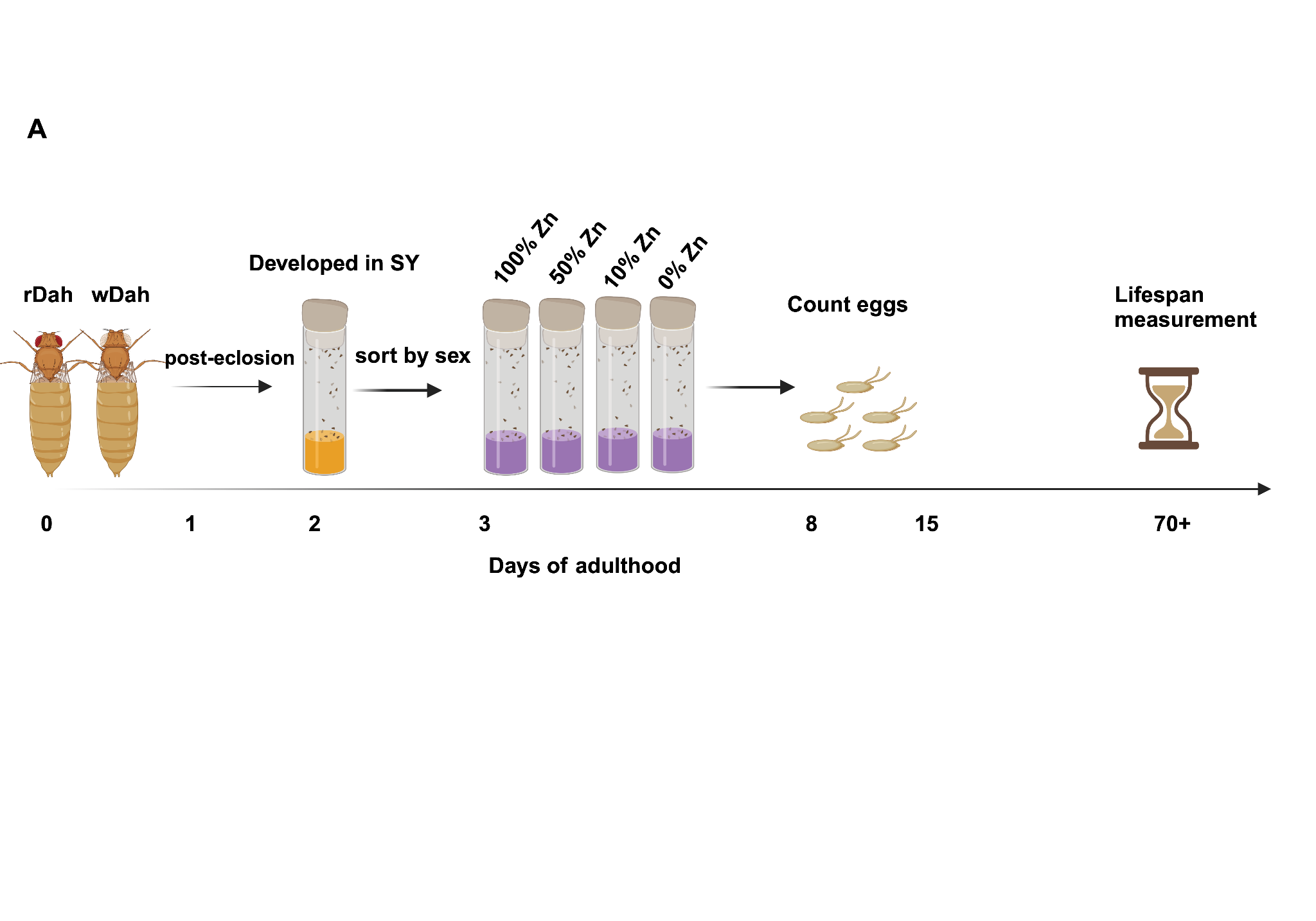

**Figure S3**

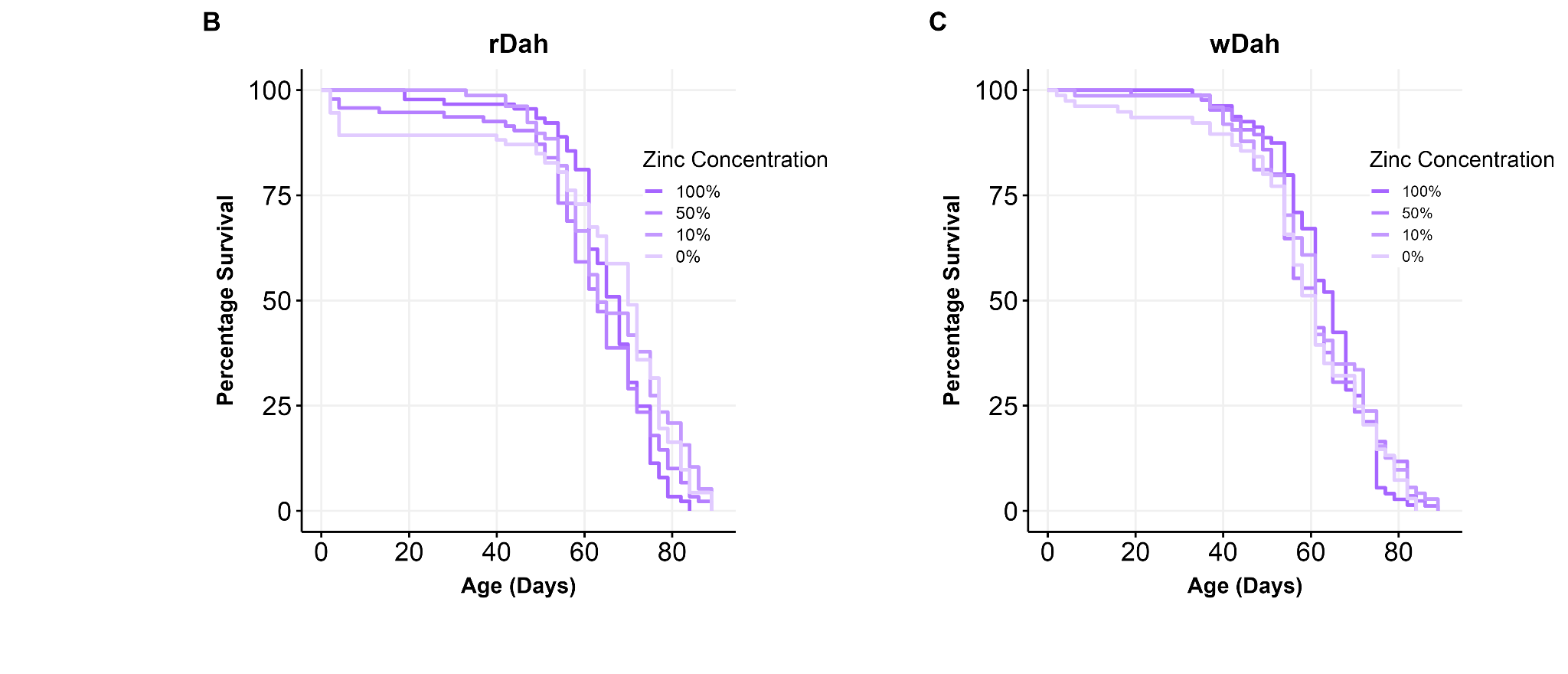

**Figure S3. Experimental timeline, dietary regimens, and lifespan analysis for adult female flies (data shown in Figure 3) in different zinc concentration diets**

*A) Flies of two genotypes, rDah and wDah were maintained separately on SY (Sugar/Yeast) food until the third day of adulthood. Once mated, on day 3, they were each transferred to four different synthetic diets. Diets included complete (100% Zinc), 50%, 10%, and 0% Zinc diets. N= 100 flies/diet*

*B) No reduction in lifespan of rDah flies fed 50%, 10% and 0% Zn diets.*

*C) There was no significant difference in lifespan amongst wDah flies fed 100%, 50%, 10% and 0% Zn diets.*

**Figure S4**

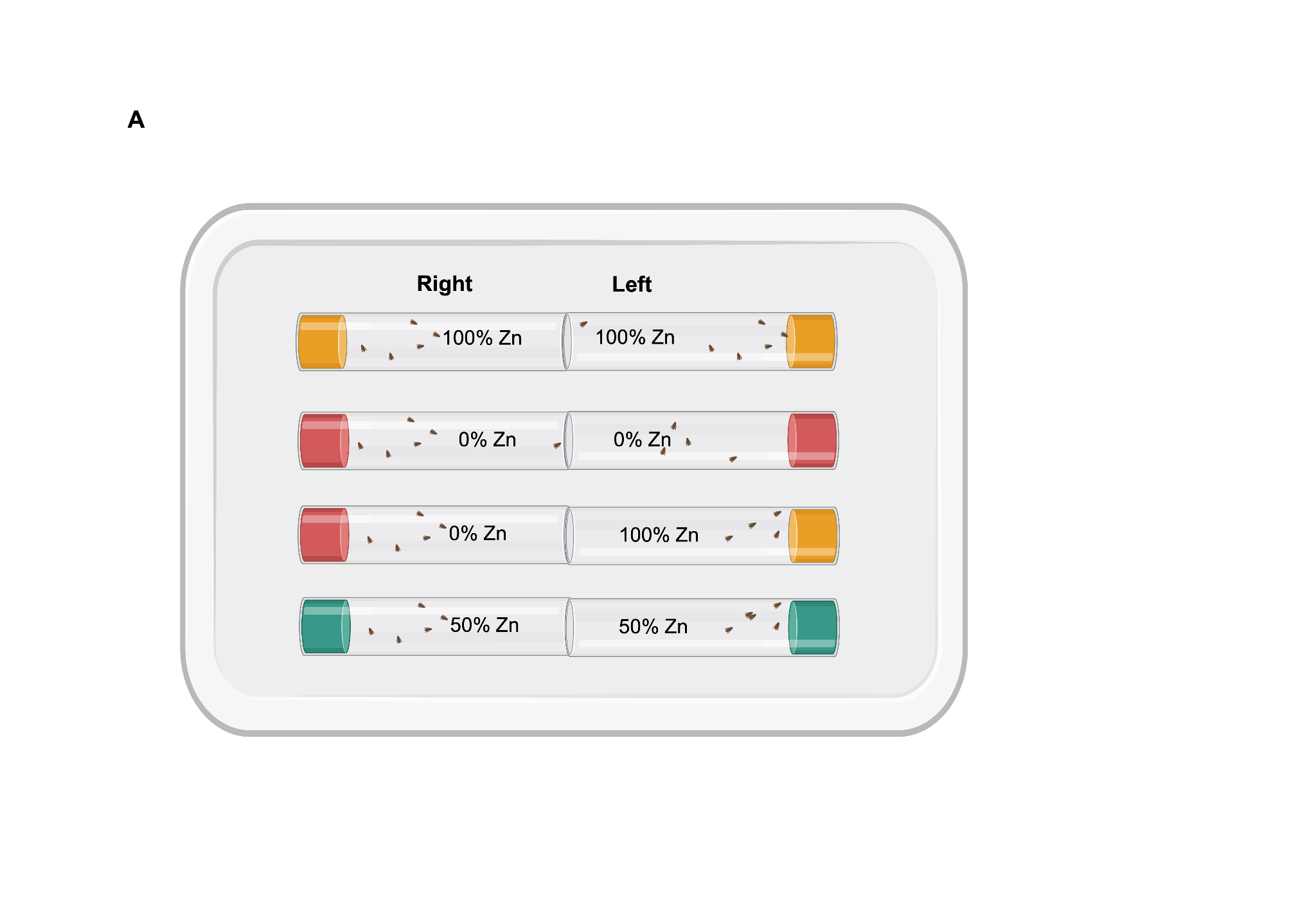

**Figure S4. Depiction of the experimental setup for measurement of feeding and egg-laying preference using the zinc choice assay for data shown in Figure 5**

*Flies were maintained on SY (Sugar/Yeast) food until the third day of adulthood. Once mated, on day 3, females were transferred to the different diet combinations: 100% Zn at both ends (100 _ 100), 0% Zn at both ends (0_ 0), 50% Zn at both ends (50_50) or vials with a choice of 0% Zn and 100% Zn (0_100). N=100 flies were used for each of the no choice diet pairs (100_100, 0 _0, 50_50) and N=150 flies were used for the choice condition (0_100).*

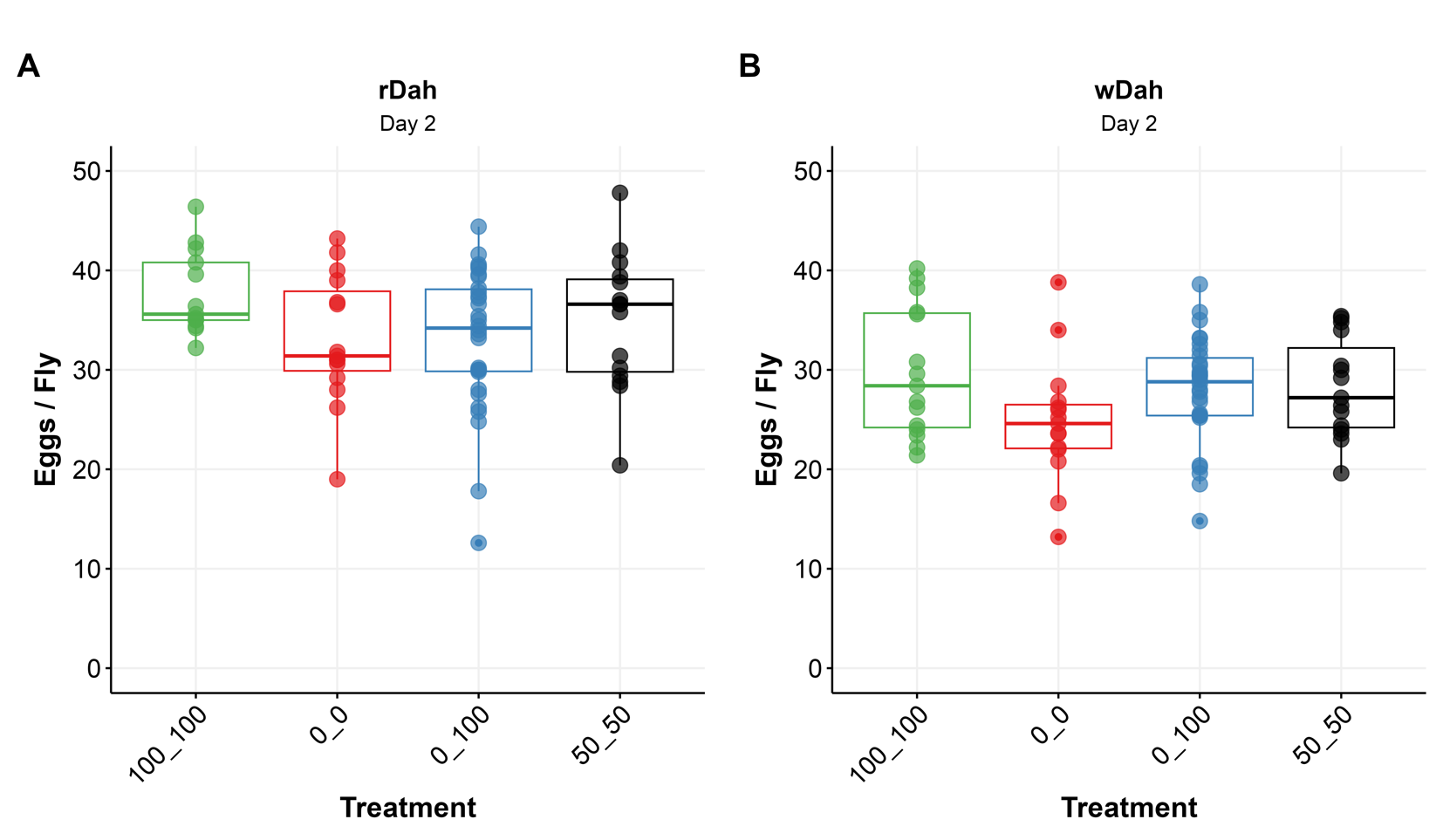

**Figure S5**

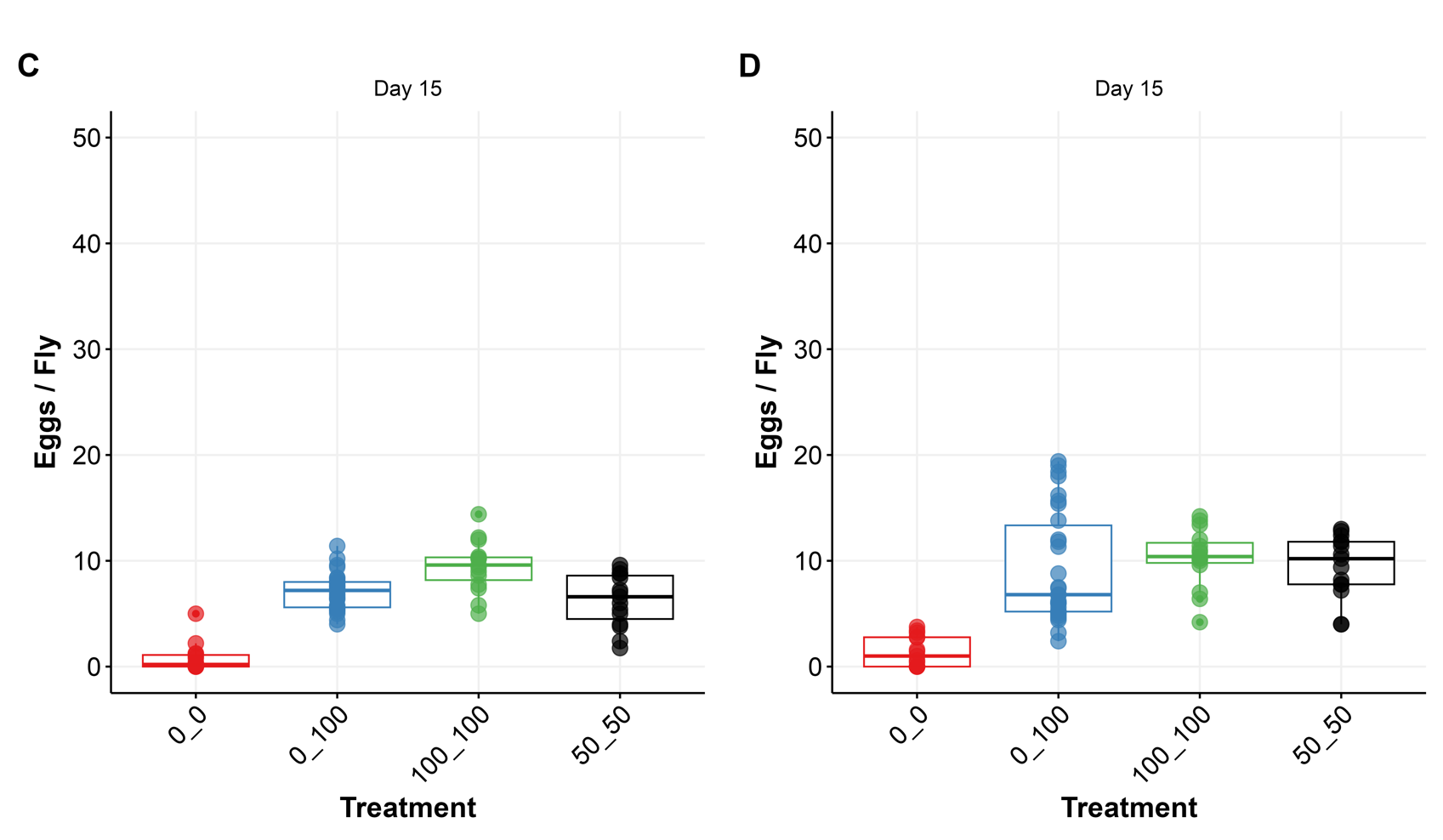

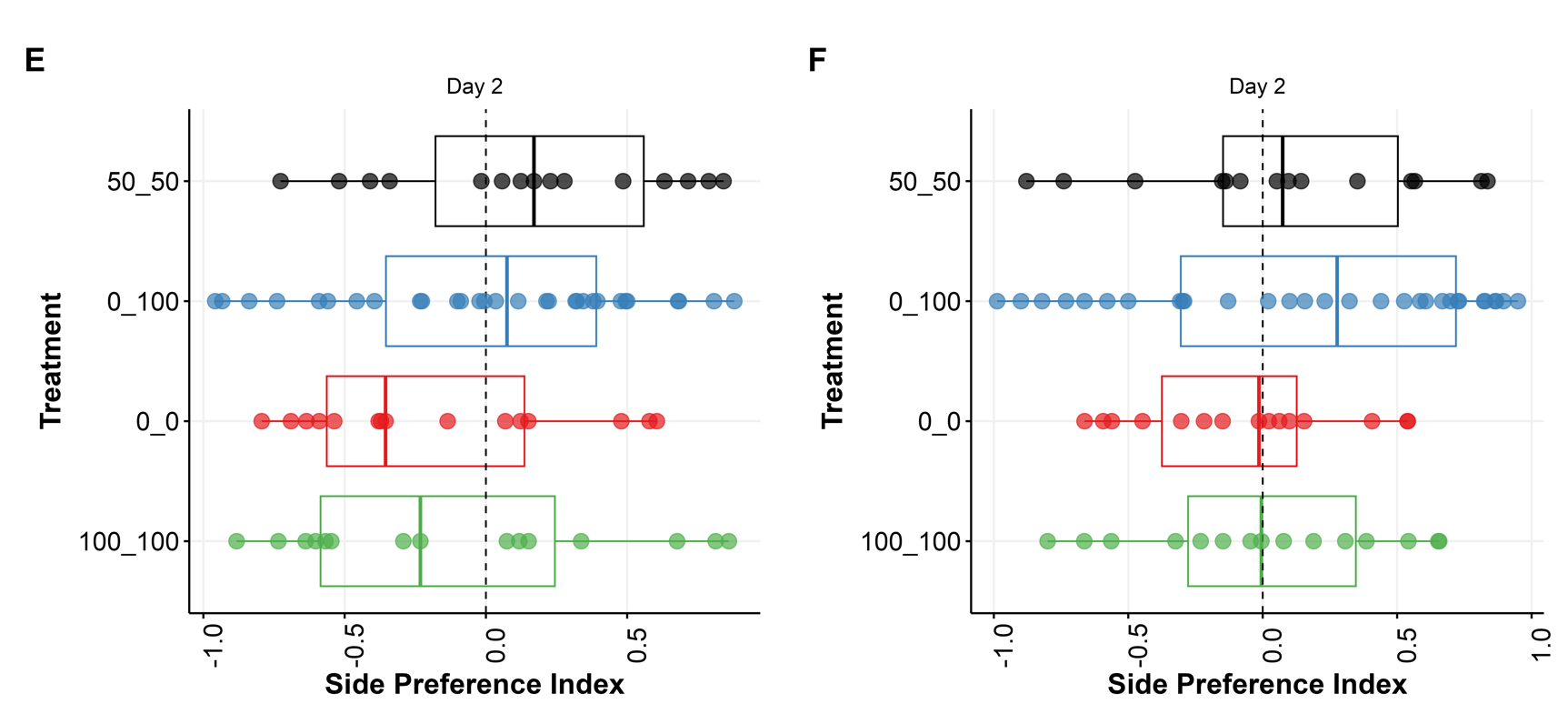

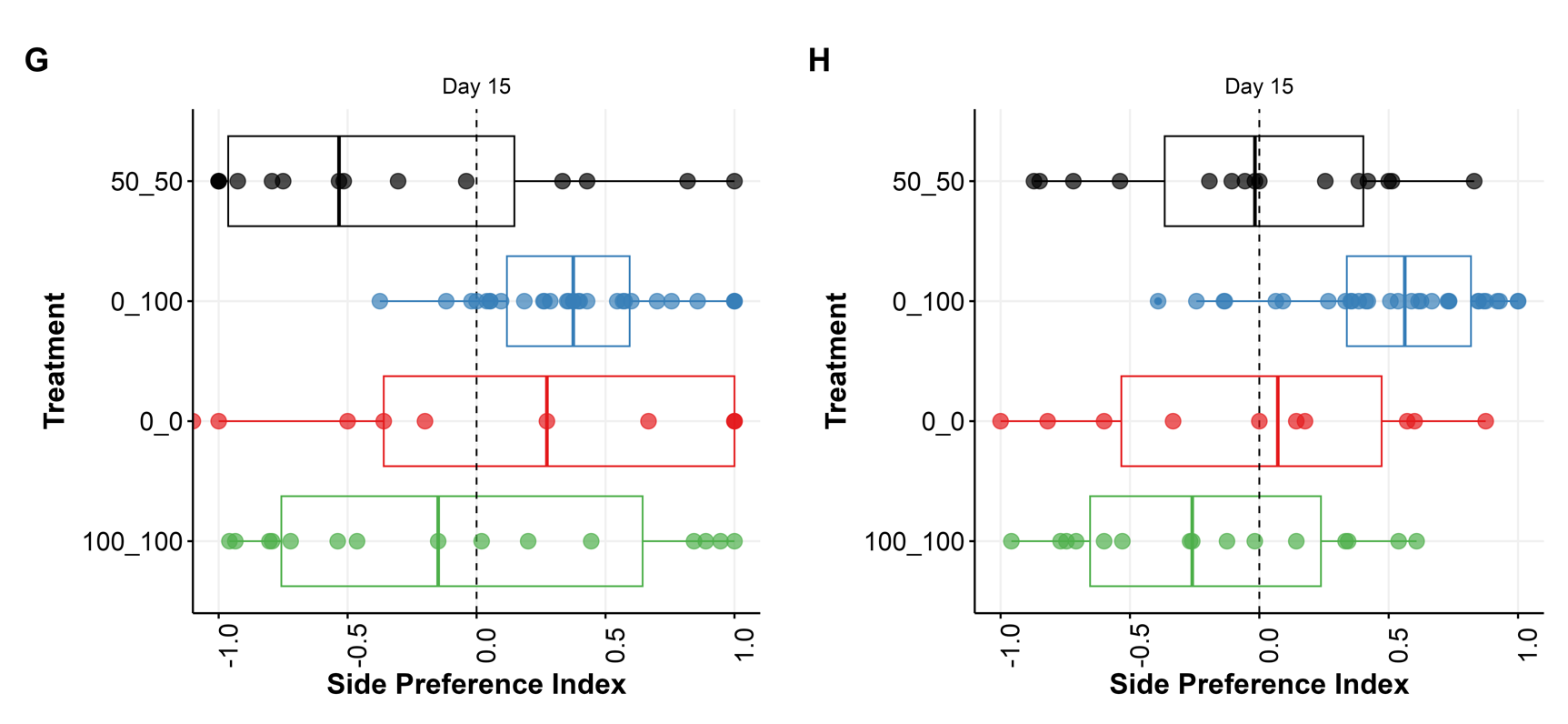

**Figure S5. Feeding behaviours on days 2 and 15, along with oviposition responses of flies on day 2, in the context of food choices varying in Zn concentration**

*A) On day 2,rDah egg-laying was almost the same across all dietary pairs with flies on the 100_100 diet combination laying a small but significantly higher number of eggs. (B) On day 2, egg-laying of wDah remained consistent across all dietary pairs. (C) On day 15, rDah and (D) wDah displayed consistent egg-laying trends across food types, resembling patterns observed on day 8. On day 2, (E) rDah showed no oviposition preference based on the Zn content of the food, while (F) wDah displayed a weak oviposition preference for food containing Zn over that with none. On day 15, (G) rDah and (H) wDah flies on day 15 showed an increase in preference to oviposit on food containing Zn over food lacking Zn.*
